# Mapping the evolutionary landscape of Zika virus infection in immunocompromised mice

**DOI:** 10.1101/839803

**Authors:** Maria G. Noval, Margarita V. Rangel, Katherine E. Johnson, Elfie De Jesus, Adam Gerber, Samantha Schuster, Ken Cadwell, Elodie Ghedin, Kenneth A. Stapleford

## Abstract

The fundamental basis of how arboviruses evolve in nature and what regulates the adaptive process remain unclear. To address this problem, we established a Zika virus (ZIKV) vector-borne transmission system in immunocompromised mice that mimics evolutionary characteristics of ZIKV infection in humans. Using this system, we defined factors that influence the evolutionary landscape of ZIKV infection and show that transmission route and specific organ microenvironments impact viral diversity and defective viral genome (DVG) production. In addition, we identified in mice the emergence of ZIKV mutants previously seen in natural infections, including variants present in currently circulating strains, as well as mutations unique to the mouse infections. With these studies, we have established an insect-to-mouse transmission model to study ZIKV evolution. We also defined how organ microenvironments and infection route impact the ZIKV evolutionary landscape, providing a deeper understanding of the factors that regulate arbovirus evolution and emergence.

## Introduction

Arboviruses (arthropod-borne viruses) exist in a complex and dynamic life cycle between an arthropod vector (e.g. mosquitoes, ticks) and a host (e.g. mammals, birds, plants) (Weaver, Charlier, Vasilakis, & Lecuit, 2017). It is within these disparate hosts that arboviruses replicate and undergo genomic evolution, which plays an essential role in transmission and pathogenesis (Parvez & Parveen, 2017; Weaver, 2006). To date, we understand little of how arboviruses evolve, are transmitted, or cause disease in nature and thus it is crucial to study these aspects of virus biology in a controlled laboratory setting.

Zika virus (ZIKV) is a positive strand RNA virus and a member of the *Flaviviridae* family (Miner & Diamond, 2017), which includes other human pathogens such as West Nile virus (WNV), dengue virus (DENV), and Yellow Fever virus. ZIKV has been responsible for several outbreaks including a recent and explosive epidemic in 2015-2016 where it swept through the Americas causing severe disease (Campos, Bandeira, & Sardi, 2015; Fauci & Morens, 2016; Lucchese & Kanduc, 2016). Importantly, although we still do not completely understand the molecular basis of ZIKV induced disease, viral genomic evolution has been implicated as a driving force for ZIKV disease and transmission in these recent epidemics (Aldunate, Gambaro, Fajardo, Sonora, & Cristina, 2017; Delatorre, Mir, & Bello, 2017; Y. Liu et al., 2017; Metsky et al., 2017; Shi et al., 2016; Yuan et al., 2017). This gap in knowledge, coupled with overall increases in the evolution of vector-borne pathogens and their spread to naïve vector and host populations (Gill, Beckham, Piquet, Tyler, & Pastula, 2019; Hermance & Thangamani, 2017), highlights the need to study arbovirus evolution and emergence during vector-borne transmission.

ZIKV can be transmitted by a variety of routes, including through sexual contact (Hastings & Fikrig, 2017) and mother-to-fetus (Coyne & Lazear, 2016), yet the primary route of infection is by the widespread and invasive mosquito species *Aedes (Ae.) aegypti and Ae. albopictus* (Magalhaes, Foy, Marques, Ebel, & Weger-Lucarelli, 2017). To date, several important studies using mouse models have addressed the physiological effects of vector-borne arbovirus transmission (Pingen et al., 2016; Pingen, Schmid, Harris, & McKimmie, 2017). These seminal studies, using human pathogens such as DENV (Cox, Mota, Sukupolvi-Petty, Diamond, & Rico-Hesse, 2012; Schmid et al., 2016), WNV (Schneider et al., 2010; Schneider et al., 2006), Semliki Forest virus (Pingen et al., 2016), chikungunya virus (CHIKV) (Agarwal et al., 2016), and Rift Valley fever virus (Le Coupanec et al., 2013) revealed that mosquito bites can enhance viral infections, increase pathology, and alter host immune responses when compared to conventional needle inoculations. However, with the exception of several studies using rhesus macaques (Aliota et al., 2018; Dudley et al., 2017), the impact of vector-borne transmission on ZIKV viral evolution and adaptation has not been extensively explored, underscoring the need to study these processes in the context of their natural route of transmission, from mosquito to mammal. In particular, the role that specific organ microenvironments play in shaping ZIKV evolution by influencing viral diversity and defective viral genomes (DVGs) in mammals has not been investigated and is essential to our understanding of how arboviruses evolve and emerge.

In this study, we allowed ZIKV infected *Ae. aegypti* mosquitoes to feed on type I interferon receptor deficient (*Ifnar ^-/-^*) mice or subcutaneously infected mice via the footpad. We then monitored disease, isolated and quantified viral RNA and infectious particles in multiple organs and performed a viral genome deep sequencing analysis to compare ZIKV diversity between the two inoculation routes. We found that while signs of disease and viral RNA accumulation do not differ between inoculation route, viral diversity and infectious viral particle production were impacted. Interestingly, viral diversity as well as the production of DVGs was organ-specific, suggesting that organ-specific selective pressures and bottlenecks may drive the emergence of new viral variants, with potential impacts on viral disease and spread. Finally, we observed the emergence of unique ZIKV variants as well as variants present in currently circulating strains. Taken together, these studies highlight the importance of transmission route and organ-specific microenvironments in regulating viral diversity and the emergence of new viral variants. These results underscore the need to understand the driving forces of viral diversity throughout the complex and complete viral life cycle of ZIKV.

## Results

### ZIKV inoculation route does not impact pathogenesis or viral RNA levels in immunocompromised mice

The study of arbovirus evolutionary trajectories within and between hosts is essential to understanding the fundamental mechanisms these viruses use for transmission and pathogenesis. ZIKV is transmitted by the *Ae.* species of mosquito; we were thus interested in studying the evolution of ZIKV during transmission from the mosquito vector to mice. We took advantage of the well-established *Ifnar^-/-^* mouse model (Lazear et al., 2016), which favors unrestricted viral replication, and the prototypical African lineage Ugandan strain of ZIKV (MR766) that has enhanced replication and pathogenesis compared to other contemporary strains of ZIKV. We hypothesized that using an evolutionary distant virus from the currently circulating strains would provide better insight into the evolutionary trajectory of ZIKV in nature. We infected *Ae. aegypti* mosquitoes via a bloodmeal containing 10^6^ plaque forming units (PFU)/ml of the infectious clone-derived MR766 virus. After an incubation period of 14 days, we allowed infected mosquitoes to feed on *Ifnar^-/-^* mice. In parallel, we infected *Ifnar^-/-^* mice subcutaneously in the footpad with 50 PFU of MR766 and compared the disease progression and viral RNA levels generated by these two different routes of inoculation (Figure 1A). We found no significant differences in weight loss (Figure 1B) and overall signs of disease (Figure 1C) between mice infected by either mosquito bite or needle-inoculation. We euthanized these mice at seven days post-infection, which coincided with the humane endpoint, and quantified viral RNA genomes from different target organs. Interestingly, we found no significant differences in viral RNA levels between inoculation routes (Figure 1D). These data are in contrast to what has been previously reported for ZIKV vector-borne infections in rhesus macaques (Dudley et al., 2017) and may suggest that in the absence of a type I interferon response, vector-borne transmission could follow a similar course of infection as seen in subcutaneous inoculations.

**Figure 1:**
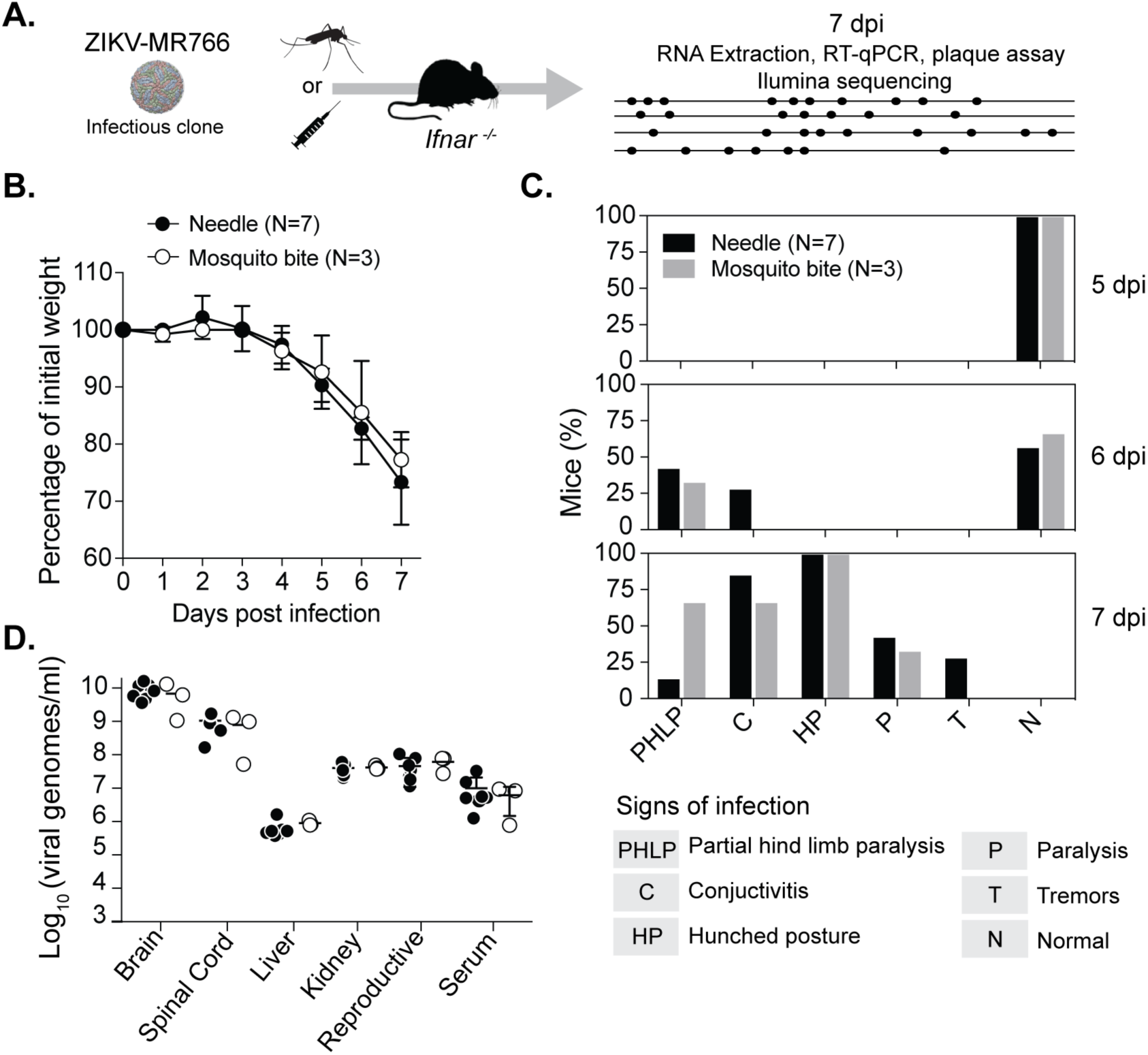
ZIKV vector-borne transmission mimics needle inoculation in disease progression and viral RNA replication. **(A).** Schematic of experimental setup. **(B).** Individual mice were infected via Ae. aegypti mosquitoes carrying ZIKV MR766 or subcutaneously via needle-inoculation with 50 PFU ZIKV MR766 in the footpad. Mice were weighed daily for seven days and euthanized at a human endpoint (20% of body weight lost). Data represent the average and SEM of three independent mosquito ¡nfections (N=3 total mice) and two independent experiments of needle inoculation (N=7 total mice). Mann-Whitney test, all data were nonsignificant (p > 0.05) **(C).** Disease was monitored during seven days post infection and the percentage of mice exhibiting signs of infection are shown. **(D).** Viral RNA genomes were quantified in each organ by RT-qPCR. Data represent the average and SEM. Mann-Whitney U test, all data were nonsignificant (p>0.05).

### ZIKV evolution in *Ifnar^-/-^* mice mimics evolutionary characteristics found in nature

Next, we asked whether the infection of these mice recapitulated the evolutionary dynamics of ZIKV observed in nature. To do this, we amplified the ZIKV protein-coding region (nt 21 to 10,528 of the ZIKV genome) in three over-lapping amplicons by PCR from: (i) our viral stock (used to infect mosquitoes for vector-borne transmission and to inoculate mice via the footpad), (ii) organs isolated from individual needle-inoculated mice, (iii) whole infected mosquitoes, and (iv) organs isolated from individual mosquito-infected mice. We successfully amplified the complete coding region from the majority of our samples and performed a deep sequencing analysis of the viral subpopulations, where we obtained similar numbers of sequencing reads for each sample (Supplemental Figure 1). Upon sequencing, we observed that our original ZIKV stock (Figure 2A, **Stock 1 - Red**) had one consensus change (variant frequency >50%) in NS5 (A9133T), the viral RNA-dependent RNA polymerase, which was also found in the plasmid and thus it was removed from further analysis (Supplemental Table 1). In addition, in stock 1 we identified 3 synonymous and 6 nonsynonymous minority variants (variant frequency between 1-50% of the population) located either in the capsid or in NS5 (Figure 2A, and Supplemental Table 1). In mice that were needle-inoculated with this stock, there was an increase in overall diversity with a number of minority variants over background (variant frequency >1%) (Figure 2B) compared to our input stock control (Figure 2A). Further, we identified two novel nonsynonymous consensus changes in Mouse 7, one in NS4B (T7582C, F2496L, 63%) and one in NS5 (G8168A, V2688M, 62.4%) (Figure 2B **and** Supplemental Table 1, **Mouse 7**).

**Figure 2:**
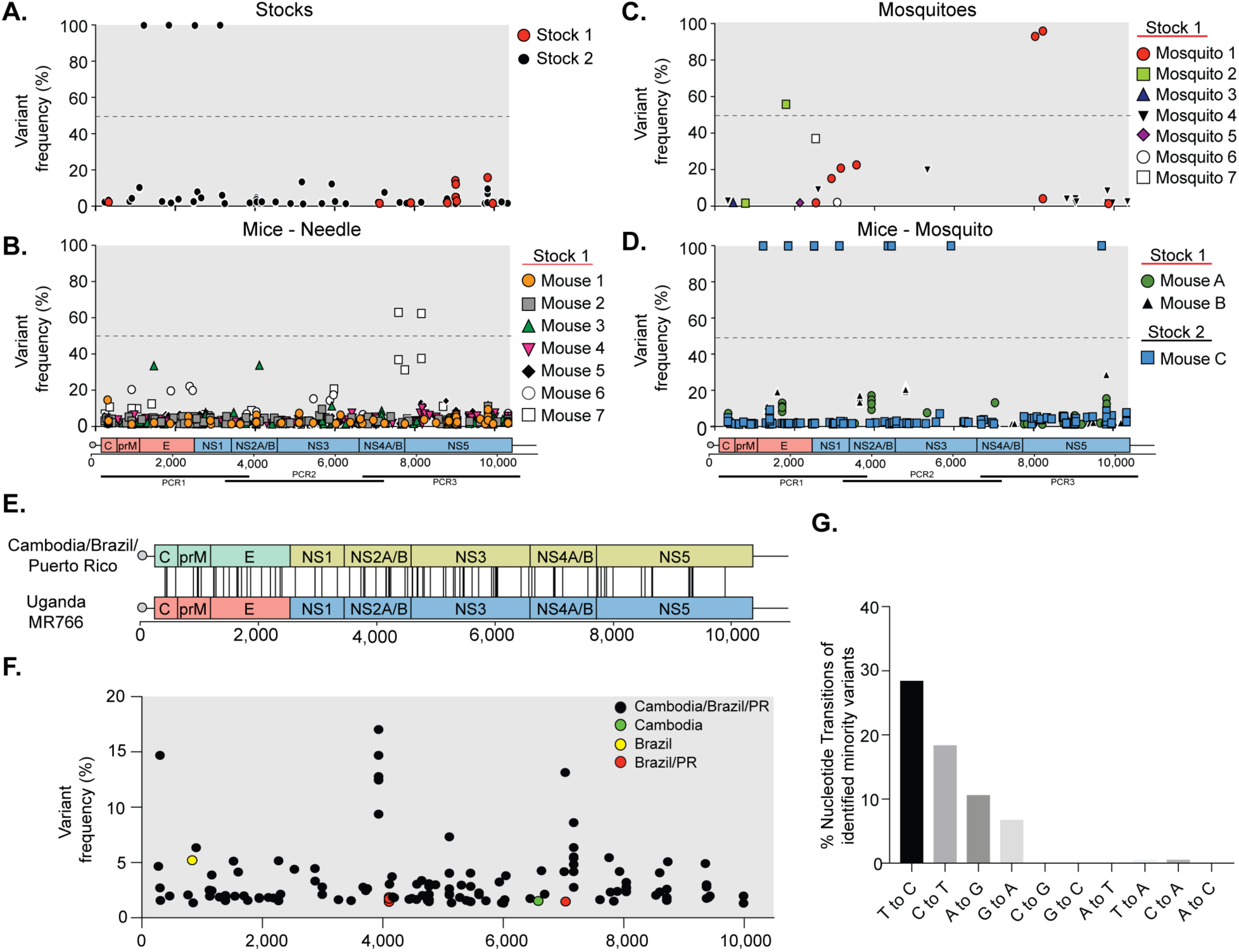
Total ZIKV single nucleotide varlant (SNV) and evolutionary analysis. The ZIKV coding region was amplified in three amplicons by PCR (bottom schematic) and each amplicon was used for deep sequencing analysis and alignment to the MR766 genome. Variants below 1% frequency were considerad background and were removed from the analysis **(A).** Viral variants present in the MR766 stocks that were used for mosquito feeding and needle infections. **(B).** Total viral variants present in mice subcutaneously infected with ZIKV MR766 (N=7). **(C).** ZIKV variants present in bodies of infected mosquitoes 14 days post feeding. After feeding on mice, mosquitoes were ground, RNA extracted, and ZIKV genome amplified by PCR. **(D).** Total viral variants present in mice infected by mosquito bite. Data represent mice from three independent experiments (N=3). **(E).** Schematic showing genomic locations of the MR766 minority variants corresponding to nucleotide changes to the Cambodian, Brazilian, and Puerto Rican strains. **(F).** Frequencies of MR766 minority variants corresponding to changes between MR766 and the Cambodian, Brazilian, and Puerto Rican strains. **(G).** Proportion of each nucleotide transition among the 86 variants identified to be different between the ZIKV strains.

In mosquitoes (Figure 2C), we found only a few minor variants above background with mosquito 1 having the majority of these mutations, including two novel consensus changes in NS5 (G8025A, G2640E, 92.9%; and C8221T, 95.9%) that were not found in the viral stock (Figure 2C **and** Supplemental Table 1). However, in mice infected via these mosquitoes (Figure 2D, **Mouse A and B**), the number of minority variants increased above background, similar to what is seen in the needle-inoculated mice. To confirm these findings with three independent infections, we repeated the vector-borne transmission with a new virus stock (Figure 2A, **Stock 2 – Black**). Upon deep sequencing of this stock, we found it to contain four synonymous consensus changes (Supplemental Table 1) and additional minority variants compared to our ZIKV plasmid and Stock 1 (Supplemental Table 2). While we were unable to obtain amplicons from mosquitoes with this infection, we observed that these variants were maintained in Mouse C, which was infected via mosquito bite from ZIKV Stock 2. Mouse C had five additional consensus changes, four of which became fixed (Supplemental Table 1). Of these new consensus changes, a nonsynonymous variant in the NS3 protein (G5962T, E1940D, 99.9%) was found at 12.1% in the viral stock. The enrichment of this variant in Mouse C could suggest that it was under positive selection in the mosquito or in the mammalian host. It could also be the result of the stochastic nature of transmission, which has tight bottlenecks. Taken together, these data suggest that ZIKV viral diversity is restricted in the mosquito host and expanded in *Ifnar ^-/-^* mice, independent of inoculation route.

Interestingly, in these analyses we observed that most minority variants were generally present at low frequencies (<30%) in mice, closely mimicking what has been seen in human isolates (Metsky et al., 2017). Moreover, we observed an increase in C to T and T to C transitions during ZIKV infection in mice, something that again was observed in humans (Metsky et al., 2017) (data not shown). Given these similarities to natural infections, we asked whether the infections in mice can recapitulate the evolutionary trajectories found in nature during the most recent outbreaks. When we compared minority variants observed in our studies with consensus differences between the Ugandan (MR766), and Cambodian, Brazilian, and Puerto Rican ZIKV strains, we found 86 unique minority variants that corresponded to exact nucleotide differences between MR766 and each of these strains (Figure 2E and F, Supplemental Table 3). Of these minority variants, 82 were common differences between MR766 and the Cambodian, Brazilian, and Puerto Rican strains, 2 were unique differences between MR766 and the Brazilian and Puerto Rican strains, 1 was a difference between MR766 and only the Brazilian strain, and 1 was a difference between MR766 and the Cambodian strain. Interestingly, these variants corresponded to 83 synonymous mutations and 3 nonsynonymous mutations with a preference for C to T and T to C transitions (Figure 2G), as seen previously in humans (Metsky et al., 2017). Finally, in addition to point mutations in currently circulating strains as compared to MR766, we also observed the emergence of 17 nonsynonymous variants that have been described as minority variants or common variants in nature (Metsky et al., 2017, Collins et al., 2019) (Supplemental Table 4). Taken together, these data suggest that ZIKV infection in *Ifnar^-/-^* can mimic various aspects of human ZIKV infections and represents a relevant model to study how the arbovirus evolves.

### ZIKV inoculation route impacts organ-specific diversity and infectious virus production

To understand how different organ microenvironments shape the genetic composition of ZIKV, we analyzed ZIKV genetic diversity in individual organs across all mice (Figure 3A). We observed that the brain, spinal cord, and liver generated more diverse populations (with single nucleotide variants above 1%) in comparison to kidney, reproductive tract, and serum. In addition, in the brain, spinal cord, and kidney there were also more high frequency variants generated in the vector-borne transmission than with needle inoculation (Figure 3A). Given this organ-specific diversity, we expected to find host and organ-specific variants during infection. Of the variants identified in Stock 1, five (at positions 9029, 9033, 9037, 9064, and 9837) were present in both the mosquito and needle-inoculated mice but were absent (or below our limits of detection) in all liver samples (Supplemental Figure 2). Further, two variants that were not present in the stock were found in both mosquito and needle-inoculated mice but were enriched (2-3 fold) in the bone marrow of the mice (NS5 C10337A, R3411S; NS5 T10340G, Y3412D). These two groups of mutations suggest that tissue-specific variants are generated within the host over the course of a ZIKV infection.

**Figure 3:**
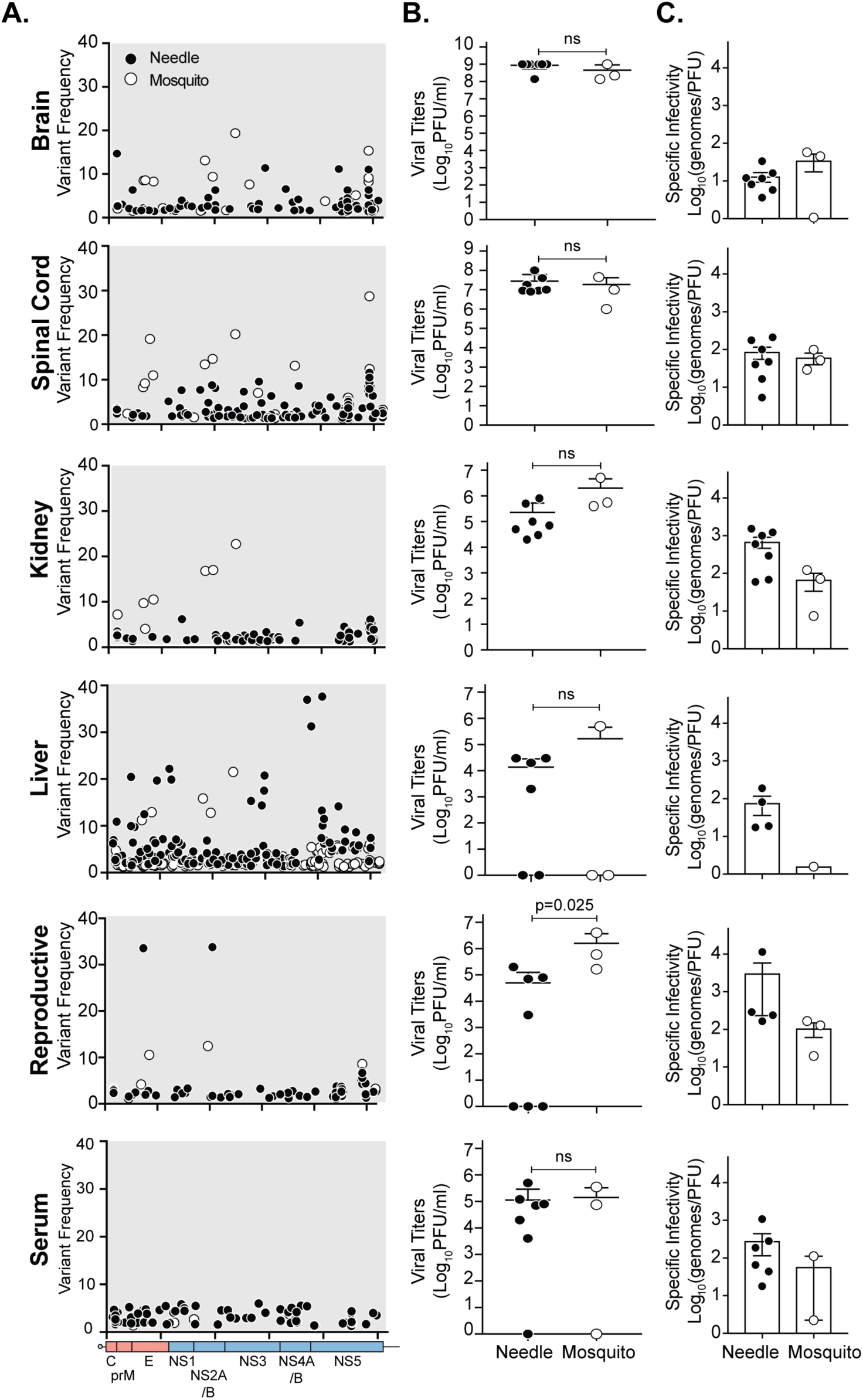
Organ-specific nucleotide variants and infectious virus particle production. **(A).** Organ-specific single nucleotide variants present in the brain, spinal cord, kidney, liver, reproductive track (ovaries and testes), and serum seven days post infection by mosquito bite (open circle) or needle inoculation (solid circle). Data represent all mice in each inoculation route. **(B).** Infectious viral titers quantified by plaque assay in each organ seven days post infection. **(C).** ZIKV Specific-infectivity in each organ and between inoculation routes. Needle inoculation (N=7) and mosquito transmitted (N=3). Data represent the average and SEM. Mann-Whitney U test, p<0.05 is considered significant, ns not significant.

We then hypothesized that although we did not see changes in viral RNA abundance in these organs, the different inoculation routes and, potentially, the different viral populations could influence infectious virus particle production. We quantified infectious ZIKV by plaque assay and found that viruses isolated from kidney and liver showed a reduced trend in the number of infectious particles generated, and the inoculation route significantly impacted the number of infectious particles isolated from the reproductive tract (Figure 3B). In line with these results, we observed organ-specific changes in viral specific-infectivity suggesting that the organ microenvironment and viral diversity contribute to ZIKV infection (Figure 3C).

Given these relative changes in organ-specific variants, we quantified both the genetic distance and the genetic diversity of each mosquito, mouse organ, and the stock sample. We focused on the sequence of PCR1, which was amplified successfully for most samples and includes the entire ZIKV structural region (Supplemental Figure 1). When calculating the pairwise genetic distance of all samples, and performing multidimensional scaling on the distance measurements, we found that samples infected with Stock 1 clustered together, while those from Stock 2 clustered independently, as expected (Figure 4A), with consensus changes and minority variants that were unique to each stock and experiment. To determine whether the trajectories of the virus were tissue specific, we focused on the degree to which each tissue sample differentiated from its respective stock virus using the calculated Euclidean distance measurements (Figure 4B). All tissues had viral populations that diversified away from their respective stock virus over the course of the infection, with the virus populations in the liver being the most dissimilar to the stock virus while the remaining tissues all maintained similar distances (Figure 4B). Further, liver samples from mice inoculated with Stock 1 virus tend to cluster more closely as observed in the MDS plots (Figure 4A), which suggests that while the virus populations in the liver are the most dissimilar to the stock virus, they evolve in the same direction, away from the virus populations in other tissues or mosquitoes (Figure 4, Supplemental Figure 3A). Together, these data suggest that even when infected with a genetically distant stock, organs shape these viral populations towards a genetic equilibrium.

**Figure 4:**
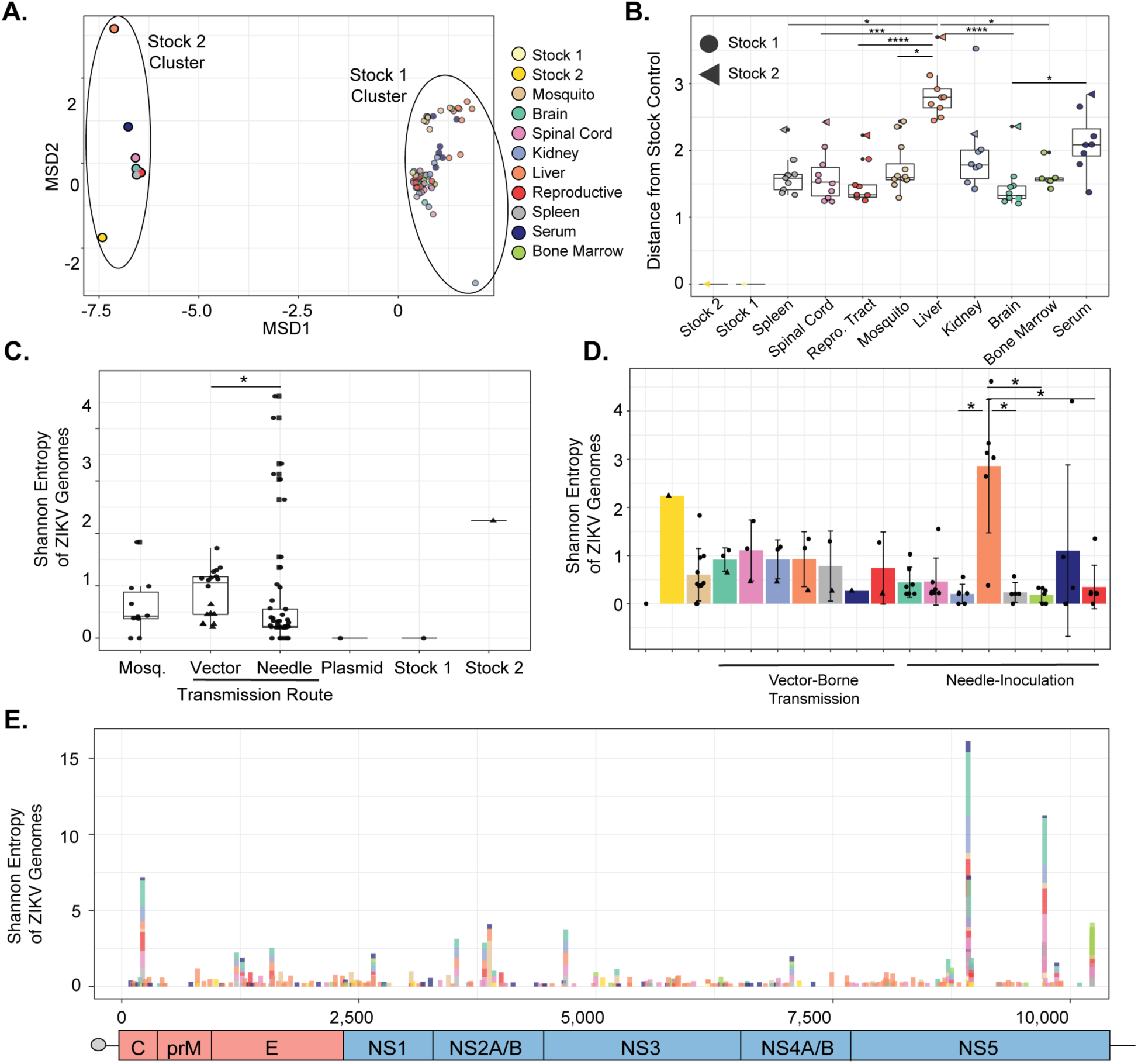
Transmission route impacts organ-specific diversity during ZIKV infection. Eucudean distance and Shannon entropy were calculated using sequence data frorri PCR1 amplicons. **(A).** Multidimensional scaling plot showing the all-versus-all Eucudean distance measurement of each organ sample. **(B).** Genetic distance of each sample from its corresponding stock control separated by tissue type. Data represent the median with inter-quartile range. Black dots represent outliers. Kruskal-Wallis with Dunn’s post-test, considering a type-I error. * p<0.05, *** p<0.001, **** p<0.0001 **(C).** Organ-specific Shannon entropy by inoculation route. Data represent the average ⊠ standard deviation. Kruskal-Wallis with Dunn’s post-test, considering a type-I error. * p<0.05 **(D).** ZIKV genetic diversity by transmission route. Data represent the median with interquartile range. Kruskal-Wallis with Dunn’s post-test, considering a type-I error. * p<0.05 **(E)** Shannon entropy calculated for each nucleotide across the complete ZIKV coding region. Color code and legend correspond to all figures.

To determine in which organs genetic diversity of the virus population was impacted the most, we quantified the Shannon Entropy from the PCR1 sequence data. The genetic diversity was affected strongly by inoculation route, with organs from mice inoculated via a needle yielding lower genetic diversity than organs from mice infected via mosquito bite (Figure 4C and D), suggesting that mosquito bite influences organ-specific viral diversity in a similar manner to what has been seen previously with rhesus macaques (Dudley et al., 2017). When we looked at each organ individually from mice infected via a needle, we found that the liver produced a highly diverse viral population followed by the brain and spinal cord (Figure 4D). In contrast, all organs from mice infected via a mosquito bite yielded relatively similar levels of genetic diversity, although higher than organs from needle-inoculated animals, with the exception of the liver (Figure 4C). This was particularly interesting as we hypothesized that Mouse C, which was infected with the high diversity Stock 2, would maintain this high diversity and be different from those inoculated with Stock 1. However, we found that organs from this mouse had lower diversity than the original stock, and had similar levels of diversity as the mice inoculated with Stock 1 (Figure 4C). These data show that independent infections with two genetic distinct inputs result in the same outcome after vector-borne transmission, suggesting that regardless of input virus diversity, the virus diversity is reshaped by either the mosquito or the mouse during vector-borne infection. Finally, when we applied these analyses to the full ZIKV coding region, we obtained similar results and identified potential diversity hot-spots across the genome (Figure 4E and Supplemental Figure 3B and C). Altogether, these data indicate that viral diversity is organ-specific and dictated (in this case) by inoculation route while still maintaining consensus sequences similar to the stock used for inoculation.

### Vector-borne transmission in *Ifnar ^-/-^* mice selects for novel ZIKV variants

To determine whether the variants generated during these ZIKV infections were potentially selected for, and thus represented potential emerging variants, we looked at the minority synonymous and nonsynonymous variants in organs from each inoculation route. We rationalized that if a virus variant was present in multiple organs of a single mouse, rather than solely in individual organs, it was likely undergoing positive selection (Figure 5, Supplemental Figure 4). In the needle-inoculated mice, we identified a number of single virus variants in individual organs (Figure 5A and B), yet while interesting these did not meet our criterion for selection. However, in the vector-borne transmission experiments we identified unique variants for two mice (Mouse A and Mouse B) that had the same point mutations in multiple organs (Figure 5C and D, **boxes**). In Mouse A, there were two synonymous changes (C1690T (8.2-12.9%) and T3931C (9.4-17.0%)). In Mouse B, there were three nonsynonymous changes (Envelope: G1316A, V114M (8.9-9.2%); NS2A: A3638G, I33V (13.1-16.8%); and NS3: G4788A, R59K (19.4-22.7%). When we mapped these changes onto the protein structures we found that the V114M mutation in the envelope was located in a highly-conserved glycerophospholipid binding pocket (Guardado-Calvo et al., 2017) (Figure 5E) and the R59K mutation in NS3 mapped to a recently identified cellular 14-3-3 binding site(Riedl et al., 2019) (Figure 5F). Finally, the I33V mutation in NS2A is located in a putative transmembrane alpha-helix predicted to be in the ER lumen (Zhang et al., 2019) (Figure 5G).

**Figure 5:**
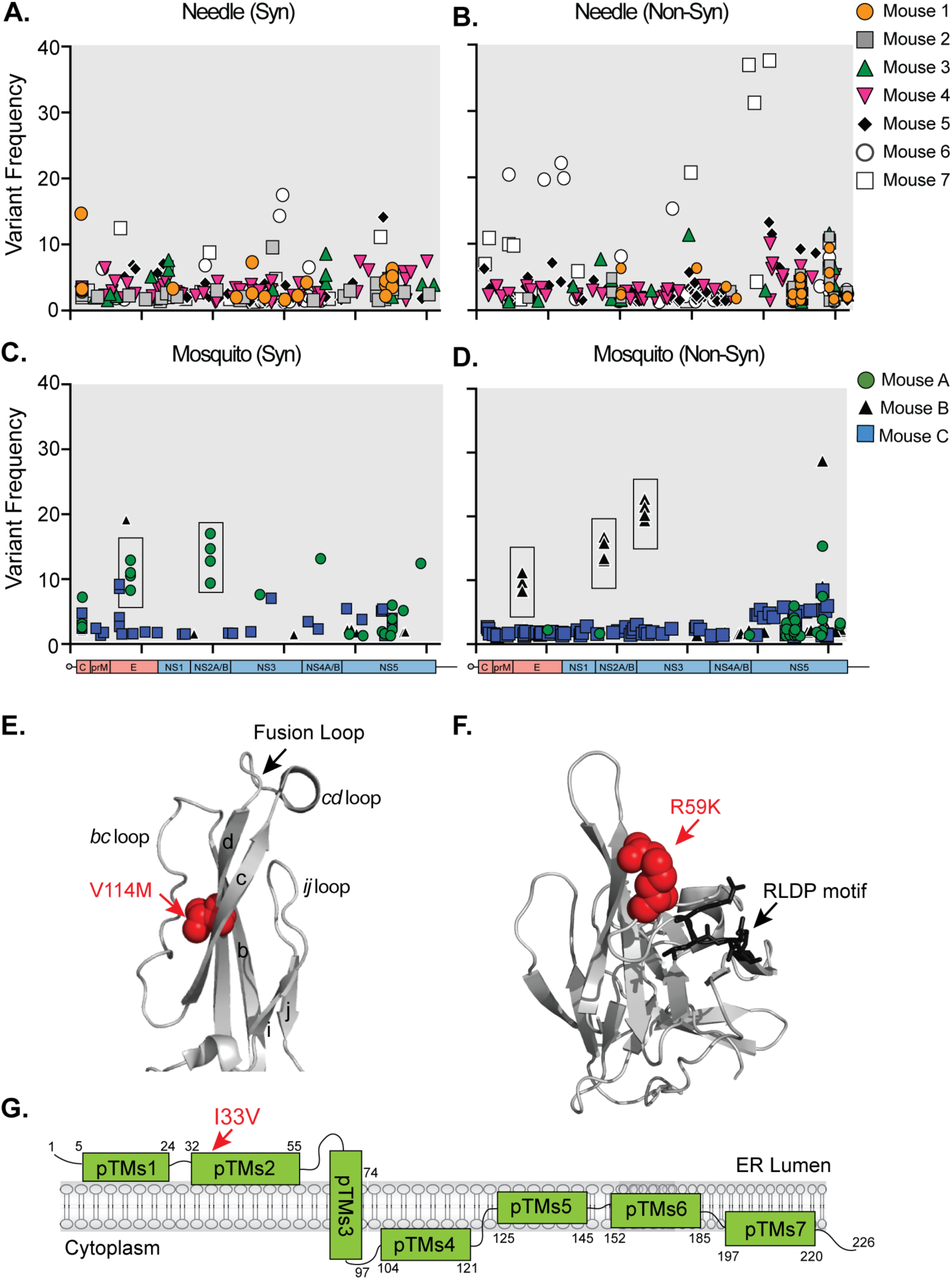
Vector-borne transmission selects for minority variants in multiple organs. ZIKV virus MR766 needle-inoculated synonymous **(A)** and nonsynonymous **(B)** and vector-transmitted synonymous **(C)** and nonsynonymous **(D)** variants present in individual mice. Boxed variants signify variants present in multiple organs of individual mice. **(E).** Structure of the tip of the ZIKV envelope protein (PDBID: 5JHM) with the V114M variant denoted in red. **(F).** Structure of the ZIKV NS3 protease (PDBID: 5GJ4) with R59K mutation in red and 14-3-3 RLDP binding motif in black. **(G).** Schematic presentation of the ZIKA NS2A protein. The I33V mutation is depicted in red.

Given the selection of these variants, we wanted to gain insight into the potential role that all identified SNVs may play in ZIKV protein function. We mapped the minority coding changes from both vector-borne transmission and needle inoculation onto the available ZIKV protein structures (Supplemental Figure 5). Mutations in the structural capsid protein localized mainly within alpha helix 4 (*α*-4), which is thought to interact with viral RNA(Shang, Song, Shi, Qi, & Gao, 2018) (Supplemental Figure 5A). In addition, variants in the nonstructural protein NS1 (Supplemental Figure 5C) are present in regions predicted to interact with the cell membrane(Brown et al., 2016), while variants in the NS3 protease (Figure 5D), and helicase (Supplemental Figure 5E) (Bukrejewska, Derewenda, Radwanska, Engel, & Derewenda, 2017), and NS5 polymerase (Zhao et al., 2017) (Supplemental Figure 5F and G) enzymes fall into key enzymatic sites. These structure-variant maps provide a framework for the future study of the molecular mechanisms underlying the function of these viral proteins, something that is poorly understood.

### Brain and spinal cord contain large numbers of DVGs during ZIKV infection

Finally, flavivirus RNA replication can lead to the production of defective viral genomes (DVGs) (Vignuzzi & Lopez, 2019; Yang et al., 2019), which can play an important role in viral interference (Brinton, 1983) and persistence (Brinton, 1982; Lancaster, Hodgetts, Mackenzie, & Urosevic, 1998; Li et al., 2011). Given this, we hypothesized that ZIKV also produces DVGs and that different organ microenvironments could influence DVG production. To address this, we mapped the coordinates of gaps within sequencing reads, corresponding to deleted regions in the genomic segments, and quantified the number of gap-spanning reads normalized to the total number of aligned sequencing reads (Figure 6). To determine if there were specific hotspots implicated in DVG generation within the ZIKV genome we looked at which genes harbored the most deletions. Interestingly, we found that capsid, pr, and M harbored the most start sites while NS1 contained predominately DVG end sites (Figure 6A and 6B). In addition, NS2A, and NS4B yielded roughly equal numbers of start and stop sites for the generation of DVGs during infection (Figure 6B). When we looked at individual DVGs in each sample, we observed that the viral input stock contained a large number of DVG gap-spanning reads (Supplemental Figure 6 and Supplemental Table 5). However, we also found DVGs present only in mice suggesting that novel DVGs are produced during infection (Figure 6C). Interestingly, different organs generated distinct DVGs during infection (Figure 6D **and** Supplemental Figure 6). In particular, the brain, spinal cord, and reproductive tract generated DVGs primarily from the PCR1 and PCR3 regions of the genome while the kidney, liver, and spleen generated roughly equal numbers of DVGs across the genome (Figure 6D). We then determined whether the transmission route impacted DVG production. While we did not find any statistical differences between the transmission routes (Supplemental Figure 7), we did observe a different abundance of DVGs in individual organs, with the brain and spinal cord having the highest number of gap-spanning reads per million, while the liver had the lowest (Figure 6D). These data are in contrast to what was seen for viral diversity (Figure 4) and may implicate DVGs as playing a role in ZIKV dissemination or pathogenesis. Taken together, these data show that organ microenvironments impact not only ZIKV genetic diversity but also the production of ZIKV DVGs, which could be acting as regulators in the viral life cycle.

**Figure 6:**
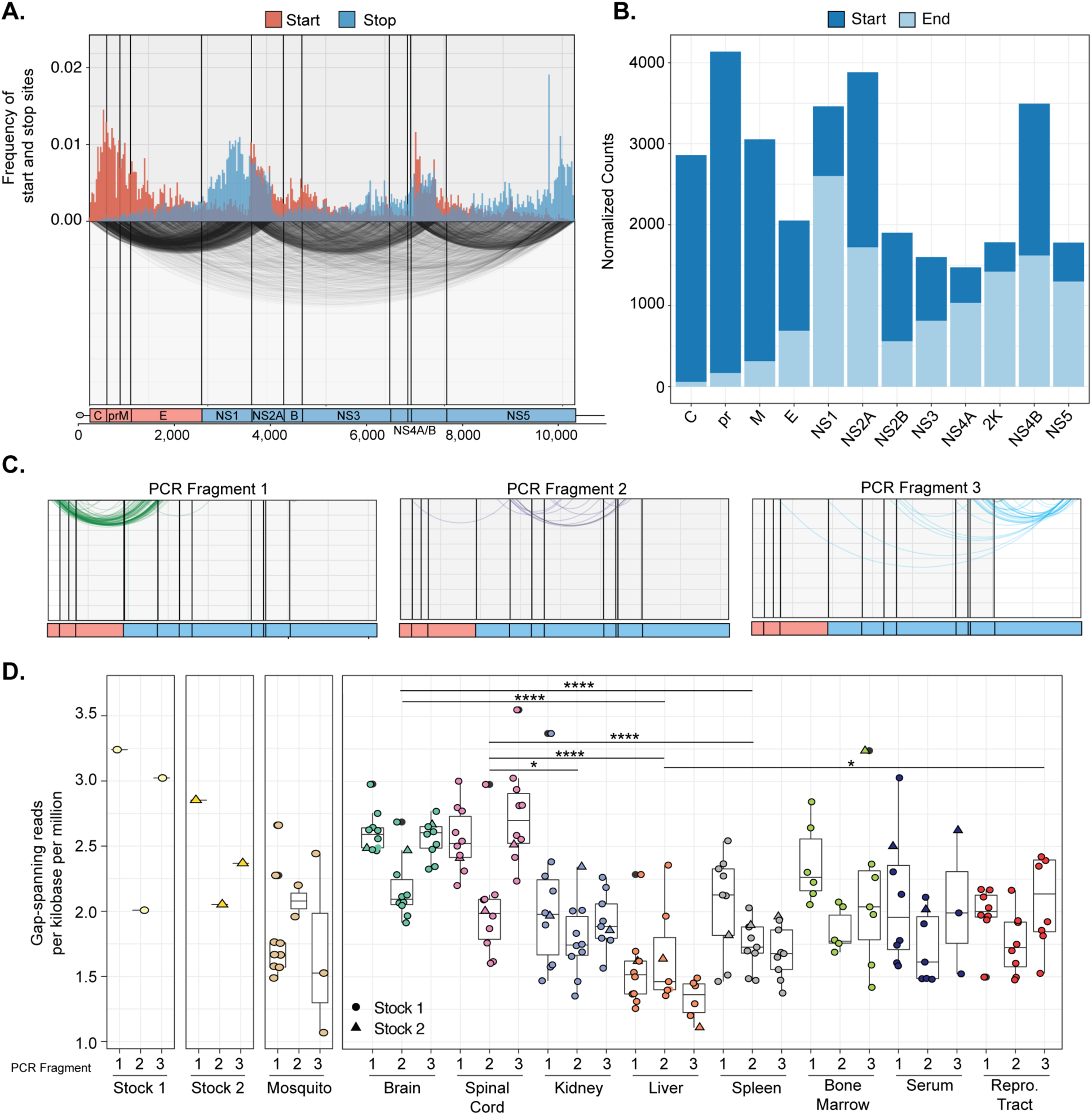
Organ-specific DVG production during ZIKV infection. **(A).** Top. Frequency of total DVG deletion start (red) and stop (blue) positions within all samples across the viral genome. Bottom. Arc plot representing the corresponding start and stop locations. **(B).** Counts of DVG species with deletion start (dark blue) and end (light blue) sites in specific ZIKV genes normalized by the gene length (kilobase). **(C).** Arc plots representing unique DVG start and stop locations found only in mice. **(D)** DVG gap-spanning reads in each organ and PCR amplicon measured by gap-spanning reads per kilobase per million. Data represent the median with interquartile range. Black dots represent outliers. Kruskal-Wallis with Dunn’s post-test, considering a type-I error. **** p<0.00001.

## Discussion

Viral infections are dynamic, involving multiple cell types, organ systems, and transmission routes. In particular, arboviruses are transmitted by insect vectors to mammalian hosts, yet we understand little of these complex infections or how vector-borne transmission influences viral diversity or emergence. In this work, we aimed to study how inoculation route and different organ microenvironments defined ZIKV pathogenesis in mice through the modulation of viral diversity, DVG production, and the emergence of new viral variants. To address this, we infected *Ifnar ^-/-^* mice with ZIKV either by the bite of a ZIKV-infected *Ae. aegypti* mosquito or by needle-inoculation. We found that the inoculation route had little impact on the course of the disease or in viral RNA levels in isolated organs. When we analyzed the viral populations present in our controls, mosquitoes, and individual mice during ZIKV infection, we thought it surprising that only a handful of high frequency (>5%) and consensus changes (>50%) could be found. These data are in contrast to what was seen in rhesus macaques where they identified a number of high frequency variants in both mosquitoes and the plasma of monkeys (Dudley et al., 2017). There are many explanations for this discrepancy, such as vector and mammalian host specificity, the viral strain used, and the use of immunocompromised animals. However, in this case it would suggest that perhaps the immune response in the monkeys would contribute to selection pressure, increasing viral diversity and leading to changes in the viral populations. It will be interesting to use immunocompetent mouse models to look specifically at the role of interferon and other host pathways on viral populations.

Nonetheless, one thing that was particularly intriguing in this study is that our data seemed to mimic human infections with the limited selection of variants, increased T to C and C to T transitions, and the majority of minority variants at around 30% of the population, or below (Metsky et al., 2017). Moreover, the MR766 African lineage of ZIKV generated minority variants that corresponded to exact nucleotide changes found in currently circulating Asian and American strains. In addition, we identified minority variants that were found in human isolates suggesting that ZIKV may be taking a similar evolutionary trajectory in mice and providing evidence that viral infections in mouse models can be used to study the evolution of the virus in human infections.

To understand how individual organs impact virus diversity and evolution, we characterized the viral populations in specific organ microenvironments. We found that each organ had distinct populations of viral variants, suggesting that viral populations may be impacted by organ and cell-specific mechanisms. This has been seen previously with multiple viruses where organ-specific bottlenecks have been studied extensively (Coffey, Forrester, Tsetsarkin, Vasilakis, & Weaver, 2013; Forrester, Guerbois, Seymour, Spratt, & Weaver, 2012; Kuss, Etheredge, & Pfeiffer, 2008; Pfeiffer & Kirkegaard, 2006; Riemersma, Steiner, Singapuri, & Coffey, 2019; Warmbrod et al., 2019; Xiao et al., 2017), supporting that this may be the case in our model, too. Interestingly, we observed high genetic diversity in the brain and spinal cord, sites of viral replication with the highest viral RNA and infectious particle loads. Moreover, the liver was an organ that generated high viral diversity, yet low viral RNA and infectious viral titers, suggesting that this diversity may be influencing viral replication. We also found that the serum, which would contain the transmittable population of virus, lacked high levels of diversity, in contrast to what was seen in monkeys (Dudley et al., 2017). However, similar to what was found by Dudley *et al*., our data support that vector-borne transmission does impact viral populations and can influence selection. The question remains as to why vector-borne transmission would specifically influence viral diversity seven days after a mosquito bite. One explanation could be that early and late inflammatory responses from the mosquito bite are exerting selection pressure impacting diversity and, as our data suggest, each individual organ may possess intrinsic mechanisms to regulate viral populations. How this diversity contributes to ZIKV fitness, pathogenesis, and transmission remains to be explored.

In addition to viral diversity, we also addressed the production of DVGs during ZIKV infection, something that, to our knowledge, had not yet been investigated. We found that the brain and spinal cord generated more DVGs – and the most diverse – as compared to other organs such as the liver, which produced the smallest number of DVGs. These data are intriguing and suggest that different organs may influence viral diversity and DVG production as a way to control the virus, or that ZIKV may take advantage of these organ-specific processes for its benefit. Interestingly, DVG production was not impacted by transmission route as it was for viral diversity. This could imply that DVG production and genetic drift are influenced by distinct factors during infection. A detailed analysis of ZIKV DVGs produced in humans and other animal models will be essential to understand the role of these truncated genomes in the viral life cycle.

We also observed the emergence of viral variants present during vector-borne transmission. In particular, three variants of interest were found in the: envelope (V114M, ∼10%), NS2A (I33V, ∼15%), and NS3 (R59K, ∼20%). The envelope variant was particularly interesting as it is located at the tip of the envelope fusion protein, near the fusion loop and the highly conserved glycerophospholipid-binding pocket. Moreover, this valine residue is located at a protein interface structurally similar to the pathogenic CHIKV variant (V80I) identified during mosquito-borne transmission (Noval, Rodriguez-Rodriguez, Rangel, & Stapleford, 2019; Stapleford et al., 2014). The structural similarities between the alphavirus E1 and flavivirus E protein are striking, and thus, we hypothesize that ZIKV V114 may play a similar role in virus biology as for CHIKV. What is not entirely clear is why this variant would only be found in one mouse and at only 10% of the viral population. One explanation could be that this mutation arose early in infection and is being lost in sick and dying mice 7 days post-infection. It will be crucial to explore the entire infectious life cycle to address this and viral evolution in future studies.

We also identified variants in the NS2A protein, which plays a role in virus assembly (Vossmann, Wieseler, Kerber, & Kummerer, 2015; Xie, Zou, Puttikhunt, Yuan, & Shi, 2015) and antagonizes the innate immune response (Chen, Wu, Wang, & Cheng, 2017; W. J. Liu et al., 2006), and another in the NS3 protease, which is close to an RLPD motif that was recently shown to be important for interactions between NS3 and cellular 14-3-3 proteins to antagonize the innate immune response (Riedl et al., 2019). The identification of these variants in immunodeficient mice is intriguing and may provide a powerful platform to identify key viral residues involved in virus-innate immune interactions. One could imagine that in immunocompromised mice, the virus no longer needs key residues to antagonize the immune response and may change these over time. It will be interesting to study the mechanistic role of the NS2A and NS3 variants in both immunocompromised and immunocompetent animal models. Finally, in these studies we have identified numerous other viral variants (Supplemental Figure 5). These mutations map to specific protein regions and potentially play key, while still unknown, roles in virus biology. Future studies addressing the fundamental mechanisms of ZIKV biology and the role the immune response and transmission (vector-borne, mother-to-child, and sexual transmission) play in driving ZIKV diversity will be essential to understanding these complex arbovirus infections.

In summary, these studies begin to define the evolutionary landscape and trajectories of ZIKV during vector-transmission and during mammalian infections. In particular, we have established an animal transmission model that can recapitulate aspects of ZIKV infection seen in humans, thus providing a powerful system to study arbovirus evolution in the lab. Using this system, we have shown that vector-borne transmission influences viral diversity and that ZIKV diversity and DVG production are impacted by the infection of various organs in immunocompromised mice. These studies of arbovirus evolution during vector-borne transmission are essential to understand the molecular mechanisms ZIKV and other arboviruses use for infection and disease, and provide a framework to study the transmission of virus variants and organ-specific forces that drive viral diversity and DVG production.

## Materials and Methods

### Cells and viruses

293T cells (ATCC CRL3216) were grown in Dulbecco’s Modified Eagle Medium (DMEM) supplemented with 1% penicillin/streptomycin (P/S), 1% nonessential amino acids, and 10% fetal bovine serum (Atlanta Biologicals) at 37°C with 5% CO_2_. Vero cells (ATCC CCL-81) were maintained in DMEM supplemented with 1% P/S and 10% newborn calf serum (Gibco) at 37°C with 5% CO_2_. All cells were verified mycoplasma free.

The Ugandan (MR766) strain of Zika virus was generated from a plasmid-based infectious clone obtained from Dr. Matthew Evans at the Ichan School of Medicine at Mt. Sinai (Schwarz et al., 2016). To generate infectious virus, 293T cells were transfected with 0.5 μg of plasmid via Lipofectamine 2000 transfection reagent (Invitrogen) and virus containing supernatants were harvested 48 hours post transfection, centrifuged at 1,200 x g for 5 mins, aliquoted and stored at −80°C. To generate a working viral stock, transfection virus stocks were used to infect Vero cells and virus containing supernatants were harvested 48 hours post infection centrifuged at 1,200 x g for 5 minutes, aliquoted, and stored at −80°C. Viral titers were determined by plaque assay on Vero cells as described below.

### Viral titrations

Viral titers were determined by plaque assay on Vero cells (Cifuentes Kottkamp et al., 2019). In brief, virus was subjected to ten-fold serial dilutions in DMEM and added to a monolayer of Vero cells for one hour at 37°C. Following incubation, a 0.8% agarose overlay was added, and cells were incubated for five days at 37°C. Five days post infection, cells were fixed with 4% formalin, the agarose overlay removed, and plaques were visualized by staining with crystal violet (10% crystal violet and 20% ethanol). Viral titers were determined on the highest dilution virus could be counted.

### Mosquito infections and manipulations

*Ae. aegypti* mosquitoes (Poza Rica, Mexico, F18-20) were a kind gift from Gregory Ebel at Colorado State University (Noval et al., 2019; Ruckert et al., 2017). Mosquitoes were reared and maintained in the NYU School of Medicine ABSL3 facility at 28°C and 70% humidity with a 12:12 hour diurnal light cycle. The day before infection, female mosquitoes were sorted and starved overnight. The day of infection, mosquitoes were exposed to an infectious bloodmeal containing freshly washed rabbit blood, 5 mM ATP, and 10^6^ PFU/ml virus for approximately 30 minutes, subsequently cold-anesthetized, and engorged female mosquitoes were sorted into new cups. Engorged mosquitoes were incubated at 28°C with 70% humidity for 14 days and fed ad libitum with 10% sucrose. Following incubation, mosquitoes were allowed to feed on naïve *Ifnar ^-/-^* mice as described below. After mouse feeding, mosquitoes were placed into a 2 ml tubes containing 200 μl phosphate buffer saline (PBS) and a steel ball. Mosquitoes were ground using a TissueLyser (Qiagen) with 30 shakes/second for 2 minutes. After grinding, each mosquito homogenate was mixed with an equal volume of Trizol (Invitrogen) for RNA extraction.

### Mouse infections and transmission studies

Animal experiments were performed in accordance with all NYU School of Medicine Institutional Animal Care and Use Committee guidelines (IACUC). Male and female *Ifnar ^-/-^* mice crossed to the C57BL/6 background were bred and housed in the animal facility at the NYU School of Medicine and subsequently transferred to the NYU School of Medicine ABSL3 animal facility for all experiments. For needle inoculations, mice were anesthetized briefly with isoflurane and inoculated in the right rear footpad with 50 PFU of ZIKV. Mice were weighed and monitored daily for signs of disease and euthanized at a defined humane endpoint where the mice had lost at least 20% of their initial body weight. Mice were dissected and organs placed in 2 ml round bottom tubes containing 500 μl of PBS and a steel ball. Organs were homogenized as described above, clarified by centrifugation at 8,000 x g for 10 minutes, and virus containing supernatants were used directly to quantify viral titers by plaque assay or mixed with equal volume of Trizol for RNA extraction and RT-qPCR.

For transmission studies (Carrau et al., 2019), ZIKV infected mosquitoes (∼5/cup) were allowed to feed on the tail of male and female *Ifnar ^-/-^* mice for 30 minutes. Following feeding, mice were placed back in their cages and the mosquitoes processed as described above. Mice were weighed and monitored daily for signs of disease and euthanized at the humane endpoint. Mice were dissected and organs were processed as described above.

### RNA extractions and RT-qPCR

Total RNA from homogenized mosquitoes and mouse organs were isolated by Trizol following the manufacturer’s instructions and resuspended in 50 μl of nuclease-free water. Total RNA was quantified, diluted to equal amounts of RNA for each organ, and viral genomes were quantified using the Taqman RNA-to-Ct kit (Applied Biosystems) using primers in Supplemental Table 6. A standard curve was generated from *in vitro* transcribed ZIKV RNA for each experiment as previously described.

### Genome amplification and library preparation

The ZIKV genome was amplified by PCR in three overlapping fragments for deep sequencing analysis. In brief, 200 ng of RNA from each sample was used to generate virus cDNA using in a Maxima H minus strand kit (Invitrogen) and ZIKV specific primers (Supplemental Table 6). cDNA was then used immediately to amplify the genome using Phusion high-fidelity DNA polymerase (Thermo) with the primers in Supplemental Table 6. PCR products were purified using a PCR cleanup kit (Macherey-Nagel) and with 0.9x AMPure beads (Beckman Culture, Inc.). One ng of purified amplicons was used as input into the Nextera DNA Library Preparation Kit (Illumina). Libraries were purified with 0.55x AMPure beads, quantified, pooled at equimolarity, and sequenced on the NextSeq500 at MidOutput 2×150 - 300 Cycle v2.

### Deep sequencing analysis

Sequencing reads were trimmed using Trimmomatic/0.36 and the Nextera adaptors file; a sliding window of 4 base pairs scanned the reads to make sure the average quality of the base pair call was above 15 and that the minimum length was at least 36 bp long. The trimmed reads from the three individual amplicons were initially aligned separately to the reference genome (MR766GenomeReference; GenBank: KX830960.1) using bowtie2 (Kim et al., 2013) --no-mixed --very-sensitive --local parameters. The alignments were then sorted with samtools (Langmead & Salzberg, 2012) and deduplicated using Picard tools MarkDuplicates. Minor variants were identified using an in-house variant caller (https://github.com/GhedinLab/ZIKV_Analysis). Coverage and minority variant calls were checked to ensure overlapping regions were identical in their nucleotide composition before merging the fastq files and then realigning the 3 amplicons to the reference file at once. Minority variants were called again on the merged alignment files and the amino acid position was added using the positions indicated on the MR766 NCBI site. Minority variants were called if coverage at the given nucleotide was at or above 500x and the frequency of the variant was above 1%, present in both forward and reverse reads, and had a quality score above 25.

### Pairwise genetic distance

Variant files generated from the first PCR alignments (PCR1) were used as input to calculate the L2-norm, which uses the Euclidean distance to perform an all-versus-all pairwise comparison of each sample at each nucleotide position.

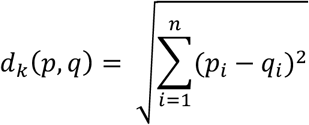

Here d_k_ is the distance between two samples at the given position *k*, where n is the total number of possible nucleotides (G,C,T,A) and p and q are the relative frequencies of the different alleles. Only frequencies of major and minor nucleotide variants were considered, all remaining nucleotides were considered to have a frequency of 0. The total distance measured between two samples, or *D*, was calculated by summing all nucleotide site distances (d_k_) across the length, *N*, of the PCR1 sequence.

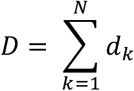

### Within-host diversity

Shannon entropy was used as the measurement for within-host diversity within each of the samples. In short, entropy scores (H) are calculated using the frequency, P_i_, for each variant at position *i* and summed across the number of alleles, S.

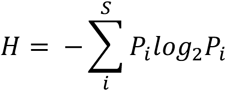

### Defective viral genome (DVG) identification

Deletion coordinates of defective viral genomes were identified by aligning the individual PCR libraries to the reference genome using the split-read aligner. Coordinates of the deletions were pulled using the CIGAR string from the alignment files (https://github.com/GhedinLab/ZIKV_Analysis). For a deletion to be called, at least 10 nucleotides had to align to the reference genome before and after the deletion. To account for noise generated during alignment start and end coordinates were clustered and grouped when located 10 nucleotides apart.

### Zika virus sequence alignments

The Ugandan (MR766, KX830960.1), Puerto Rican (PRVABC59, MK713748.1), Brazilian (Paraiba, KX280026.1), and Cambodian (FSS13025, MH158236.1) ZIKV genome sequences were aligned using MegAlign (www.dnastar.com).

### Protein representations

All protein representations were performed using Pymol (https://pymol.org/).

### Data availability

Sequencing data that support the findings of this study have been deposited in the Sequence Read Archive (SRA) under BioProject ID PRJNA589089.

### Statistics

All statistical analyses were performed using GraphPad Prism and R Studio. Data represent three independent mosquito feeds and three independent mouse infections (N=3 total mice) and two independent needle infections (N=7 total mice). Data are represented as the average ± the standard error of the mean (SEM). Mann-Whitney U and Kruskal-Wallis with Dunn’s post-test, considering a type-I error were performed and indicated in the figure legends. P values >0.05 were considered nonsignificant (ns).

## Acknowledgments.

We would like to thank all members of the Stapleford Lab for helpful comments on this manuscript. is supported by a Jan Vilcek/David Goldfarb Fellowship from the New York University Department of Microbiology, New York University. M.V.R. and K.E.J are supported in part by the Public Health Service Institutional Research Training Award T32 AI007180.

---

**Supplemental Table 1:** ZIKV MR766 consensus changes identified in this study.

| Sample | Infection Route | Nucleotide Change | Frequency | ZIKV Protein | Amino Acid Substitution in the polyprotein |
| --- | --- | --- | --- | --- | --- |
| Stock 2 |  | G1213A | 99.9% | E |  |
| Stock 2 |  | C1840T | 99.9% | E |  |
| Stock 2 |  | T2491C | 99.9% | NS1 |  |
| Stock 2 |  | A3127G | 99.9% | NS1 |  |
| Mouse 7 | Needle | T7592C | 63.0% | NS4B | F2496L |
| Mouse 7 | Needle | G8168A | 62.4% | NS5 | V2688M |
| Mosquito 1 |  | G8025A | 92.9% | NS5 | G2640E |
| Mosquito 1 |  | C8221T | 95.9% | NS5 |  |
| Mosquito 2 |  | C1783A | 55.7% | E |  |
| Mosquito 7 |  | A9133T | 99.9% | NS5 |  |
| Mouse C | Mosquito | G1213A | 99.9% | E |  |
| Mouse C | Mosquito | C1840T | 99.9% | E |  |
| Mouse C | Mosquito | T2491C | 99.9% | NS1 |  |
| Mouse C | Mosquito | A3127G | 99.9% | NS1 |  |
| Mouse C | Mosquito | C4342T | 99.9% | NS2B |  |
| Mouse C | Mosquito | T4435C | 99.9% | NS2B |  |
| Mouse C | Mosquito | G5926T | 99.9% | NS3 | E1940D |
| Mouse C | Mosquito | T9718C | 99.9% | NS5 |  |

**Supplemental Table 2:** Minority variants found in viral stocks (>1% frequency)

| Sample | Nucleotide Change | Minority Allele Frequency | ZIKV Protein | Amino Acid Substitution in the polyprotein |
| --- | --- | --- | --- | --- |
| Stock 1 | G325A | 2.3% | Capsid |  |
| Stock 1 | A7909G | 1.8% | NS5 |  |
| Stock 1 | A8820T | 2.3% | NS5 | E2905V |
| Stock 1 | A9029G | 14% | NS5 | M2975V |
| Stock 1 | T9033G | 4.9% | NS5 | M2976R |
| Stock 1 | A9037G | 12.1% | NS5 |  |
| Stock 1 | C9064G | 2.8% | NS5 | F2986L |
| Stock 1 | G9837A | 15.7% | NS5 | R3244K |
| Stock 1 | C9965T | 1.5% | NS5 | L3287F |
| Stock 2 | T229C | 2.0% | Capsid |  |
| Stock 2 | G325A | 2.6% | Capsid |  |
| Stock 2 | C853T | 2.8% | M |  |
| Stock 2 | G910A | 4.4% | M |  |
| Stock 2 | T1099C | 9.7% | E |  |
| Stock 2 | C1669T | 2.6% | E |  |
| Stock 2 | C1894T | 1.8% | E |  |
| Stock 2 | C2080T | 3.3% | E |  |
| Stock 2 | G2461A | 2.1% | E | M785I |
| Stock 2 | T2554C | 7.6% | NS1 |  |
| Stock 2 | C2650T | 4.6% | NS1 |  |
| Stock 2 | C3175T | 6.3% | NS1 |  |
| Stock 2 | T3240C | 1.4% | NS1 | L1045P |
| Stock 2 | C3869T | 1.7% | NS2A |  |
| Stock 2 | C3973T | 1.6% | NS2A |  |
| Stock 2 | G4032C | 4.7% | NS2A | R1309P |
| Stock 2 | C4045T | 2.5% | NS2A |  |
| Stock 2 | A4171G | 2.3% | NS2A |  |
| Stock 2 | C4519T | 2.0% | NS2B |  |
| Stock 2 | C4891T | 1.3% | NS3 |  |
| Stock 2 | C5177T | 13.4% | NS3 |  |
| Stock 2 | A5209G | 1.3% | NS3 |  |
| Stock 2 | C5679T | 1.8% | NS3 | S1858F |
| Stock 2 | G5926T | 12.1% | NS3 | E1940D |
| Stock 2 | A6009G | 2.9% | NS3 | K1968R |
| Stock 2 | T6277A | 1.2% | NS3 |  |
| Stock 2 | T6278G | 1.4% | NS3 | Y556D |
| Stock 2 | A6279G | 1.4% | NS3 | Y556C |
| Stock 2 | C7090T | 2.1% | NS4B |  |
| Stock 2 | T7248C | 7.4% | NS4B |  |
| Stock 2 | T7630C | 1.7% | NS4B |  |
| Stock 2 | T7804C | 1.6% | NS5 |  |
| Stock 2 | G8171T | 2.1% | NS5 | G2689W |
| Stock 2 | A8820T | 4.0% | NS5 | E2905V |
| Stock 2 | G8827A | 1.5% | NS5 |  |
| Stock 2 | A9029G | 3.6% | NS5 | M2975V |
| Stock 2 | T9033G | 1.4% | NS5 | M2976R |
| Stock 2 | G9036A | 1.4% | NS5 | G2977E |
| Stock 2 | A9037G | 3.5% | NS5 |  |
| Stock 2 | C9064G | 5.3% | NS5 | F2986L |
| Stock 2 | A9769G | 1.6% | NS5 |  |
| Stock 2 | A9831T | 1.6% | NS5 | D3242V |
| Stock 2 | T9832G | 1.8% | NS5 | D3242E |
| Stock 2 | G9837A | 9.5% | NS5 | R3244K |
| Stock 2 | C9965T | 2.2% | NS5 | L3287F |
| Stock 2 | C10187T | 2.3% | NS5 |  |
| Stock 2 | C10339T | 1.6% | NS5 |  |

**Supplemental Table 3:** MR766 minority variants present in currently circulating ZIKV strains.

| **Nucleotide Change** | **Minority Allele Frequency** | **ZIKV Protein** | **Amino Acid Substitution in the polyprotein** |
| --- | --- | --- | --- |
| T274C | 4.67% | C |  |
| T301C | 14.68% | C |  |
| C307T | 1.57% | C |  |
| T460C | 1.98% | C |  |
| T766C | 2.07% | M |  |
| **C835T<sup>a</sup>** | **5.21%<sup>a</sup>** | **M<sup>a</sup>** |  |
| C832T | 1.36% | M |  |
| C901T | 6.34% | M |  |
| G1120A | 2.50% | E |  |
| T1129C | 1.96% | E |  |
| C1159T | 3.88% | E |  |
| T1288A | 1.88% | E |  |
| T1387C | 2.02% | E |  |
| C1522T | 5.13% | E |  |
| C1543T | 1.96% | E |  |
| T1600C | 4.14% | E |  |
| G1696A | 1.99% | E |  |
| G1759A | 1.58% | E |  |
| A1948G | 1.76% | E |  |
| T2102C | 1.84% | E |  |
| T2188C | 1.79% | E |  |
| C2269T | 5.15% | E |  |
| T2302C | 2.15% | E |  |
| T2533C | 4.40% | NS1 |  |
| C2875A | 4.46% | NS1 |  |
| C2878T | 3.33% | NS1 |  |
| T2995C | 2.10% | NS1 |  |
| T3265C | 1.65% | NS1 |  |
| C3475T | 1.56% | NS1 |  |
| C3673T | 2.77% | NS2A |  |
| T3717C | 2.46% | NS2A | V1204A |
| A3742G | 2.65% | NS2A |  |
| T3931C | 17.03% | NS2A |  |
| **G4105A<sup>b</sup>** | **1.87%<sup>b</sup>** | **NS2A<sup>b</sup>** |  |
| T4111C | 3.04% | NS2A |  |
| G4153A | 3.72% | NS2A |  |
| A4168G | 1.88% | NS2A |  |
| A4189G | 1.83% | NS2A |  |
| T4438C | 1.68% | NS2B |  |
| A4483G | 1.75% | NS2B |  |
| T4561C | 1.66% | NS2B |  |
| T4567C | 3.04% | NS2B |  |
| A4645G | 3.38% | NS3 |  |
| A4660G | 2.02% | NS3 |  |
| A4744G | 1.57% | NS3 |  |
| T4768C | 1.59% | NS3 |  |
| C4875T | 3.00% | NS3 | A1590V |
| T5104C | 4.03% | NS3 |  |
| A5143G | 1.35% | NS3 |  |
| G5278A | 1.84% | NS3 |  |
| G5305A | 2.97% | NS3 |  |
| T5389C | 1.58% | NS3 |  |
| T5443C | 1.50% | NS3 |  |
| C5458T | 4.04% | NS3 |  |
| C5461T | 2.85% | NS3 |  |
| T5599C | 2.26% | NS3 |  |
| A5713G | 1.54% | NS3 |  |
| C5935T | 3.23% | NS3 |  |
| T5974C | 1.36% | NS3 |  |
| T6004C | 1.32% | NS3 |  |
| C6025T | 1.64% | NS3 |  |
| T6043C | 3.81% | NS3 |  |
| C6448T | 1.74% | NS3 |  |
| **T6583C<sup>c</sup>** | **1.53%<sup>c</sup>** | **NS4A<sup>c</sup>** |  |
| C6634T | 4.26% | NS4A |  |
| T6688C | 2.13% | NS4A |  |
| T7012C | 4.18% | NS4B |  |
| C7030T | 13.15% | NS4B |  |
| **T7036C<sup>b</sup>** | **1.48%<sup>b</sup>** | **NS4B<sup>b</sup>** |  |
| T7171C | 5.50% | NS4B |  |
| C7607T | 1.65% | NS4B |  |
| T7759C | 5.44% | NS5 |  |
| G7780A | 2.33% | NS5 |  |
| G7837A | 1.50% | NS5 |  |
| T7897C | 2.40% | NS5 |  |
| C7930T | 4.20% | NS5 |  |
| C8044T | 3.20% | NS5 |  |
| A8536G | 2.05% | NS5 |  |
| A8599G | 4.07% | NS5 |  |
| C8716T | 1.83% | NS5 |  |
| C8731T | 5.08% | NS5 |  |
| A9356G | 4.91% | NS5 | T3084A |
| T9376C | 2.78% | NS5 |  |
| T9409C | 1.84% | NS5 |  |
| A9430G | 1.98% | NS5 |  |
| T9988C | 1.33% | NS5 |  |

Bold denotes unique variants found between MR766 and Brazilian/Puerto Rican/Cambodian strains. <sup>a</sup>Different between MR766 and Brazilian strain; <sup>b</sup>Different between MR766 and Brazil/Puerto Rico; <sup>c</sup>Different between MR766 and Cambodian strain.

**Supplemental Table 4:**
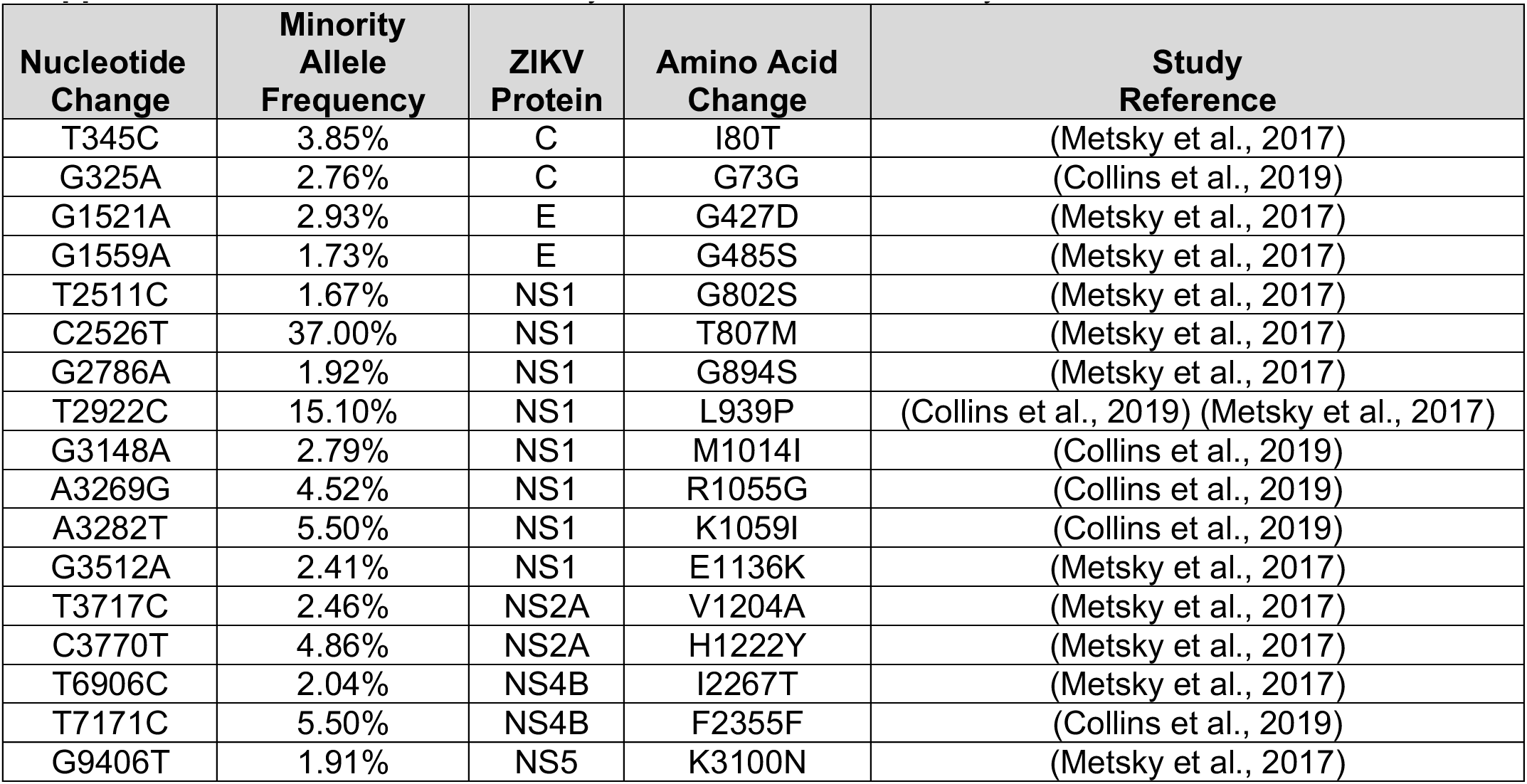
Natural minority variants found in this study.

| **Nucleotide Change** | **Minority Allele Frequency** | **ZIKV Protein** | **Amino Acid Change** | **Study Reference** |
| --- | --- | --- | --- | --- |
| T345C | 3.85% | C | I80T | (Metsky et al., 2017) |
| G325A | 2.76% | C | G73G | (Collins et al., 2019) |
| G1521A | 2.93% | E | G427D | (Metsky et al., 2017) |
| G1559A | 1.73% | E | G485S | (Metsky et al., 2017) |
| T2511C | 1.67% | NS1 | G802S | (Metsky et al., 2017) |
| C2526T | 37.00% | NS1 | T807M | (Metsky et al., 2017) |
| G2786A | 1.92% | NS1 | G894S | (Metsky et al., 2017) |
| T2922C | 15.10% | NS1 | L939P | (Collins et al., 2019) (Metsky et al., 2017) |
| G3148A | 2.79% | NS1 | M1014I | (Collins et al., 2019) |
| A3269G | 4.52% | NS1 | R1055G | (Collins et al., 2019) |
| A3282T | 5.50% | NS1 | K1059I | (Collins et al., 2019) |
| G3512A | 2.41% | NS1 | E1136K | (Metsky et al., 2017) |
| T3717C | 2.46% | NS2A | V1204A | (Metsky et al., 2017) |
| C3770T | 4.86% | NS2A | H1222Y | (Metsky et al., 2017) |
| T6906C | 2.04% | NS4B | I2267T | (Metsky et al., 2017) |
| T7171C | 5.50% | NS4B | F2355F | (Collins et al., 2019) |
| G9406T | 1.91% | NS5 | K3100N | (Metsky et al., 2017) |

**Supplemental Table 5:**
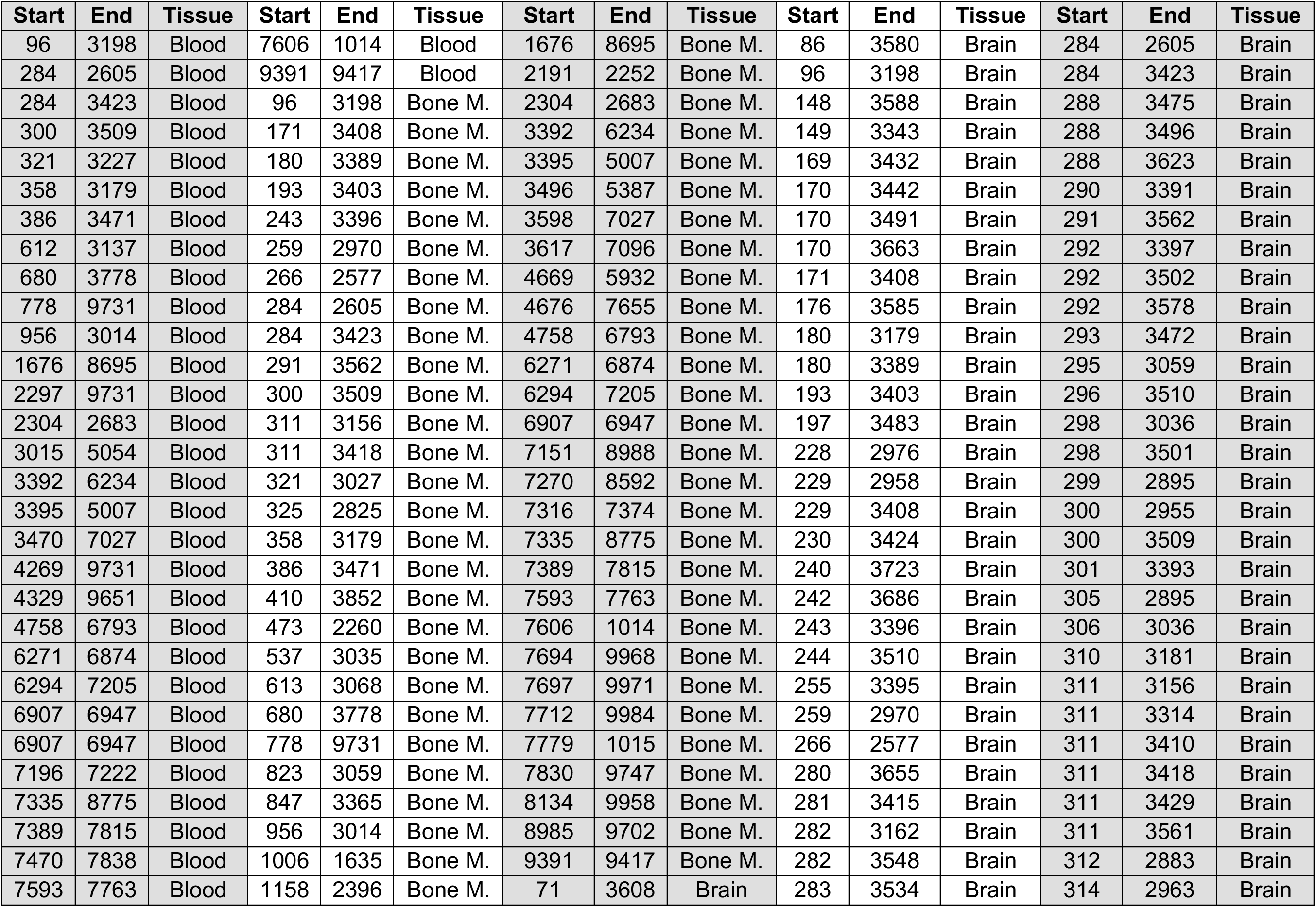
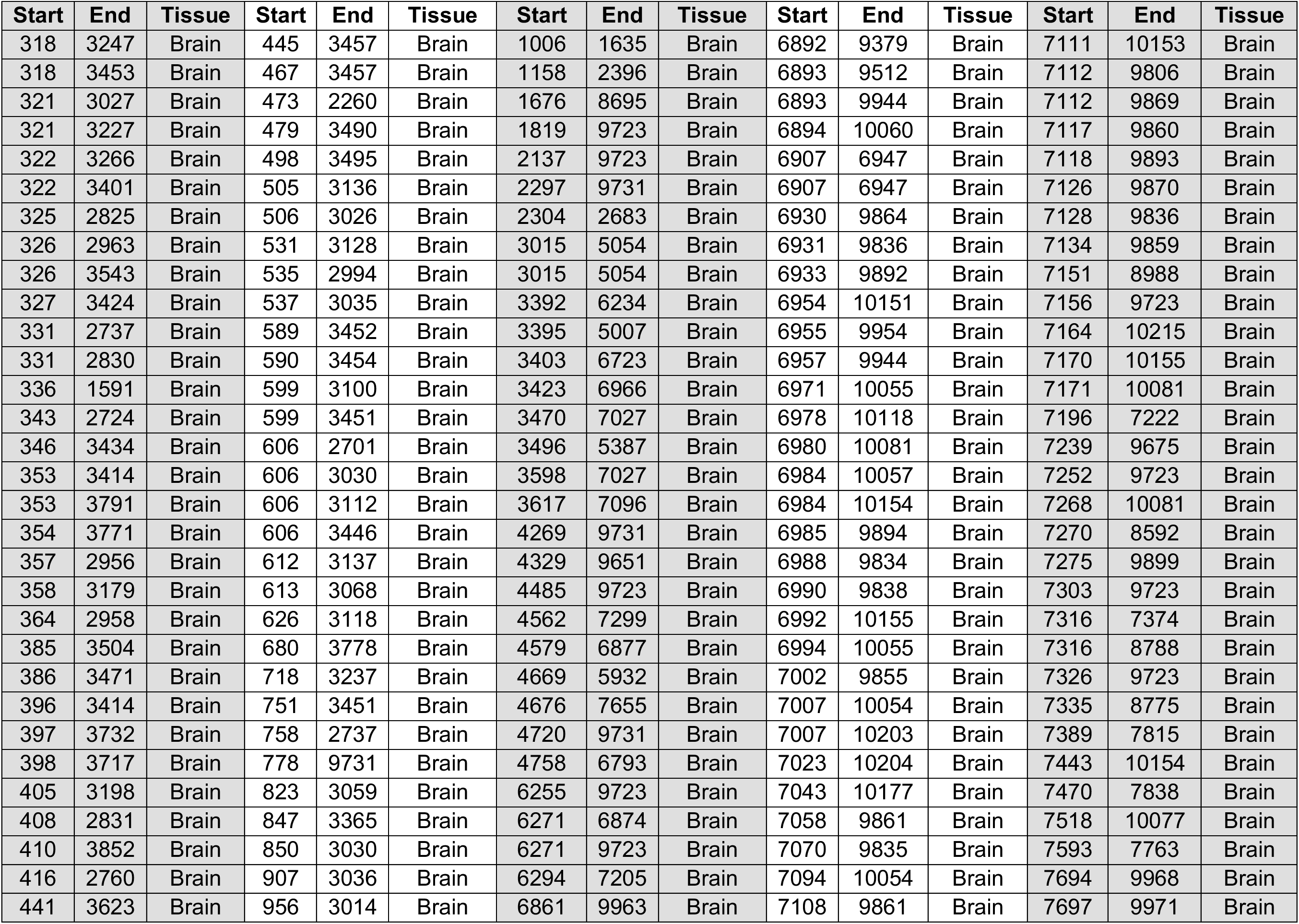
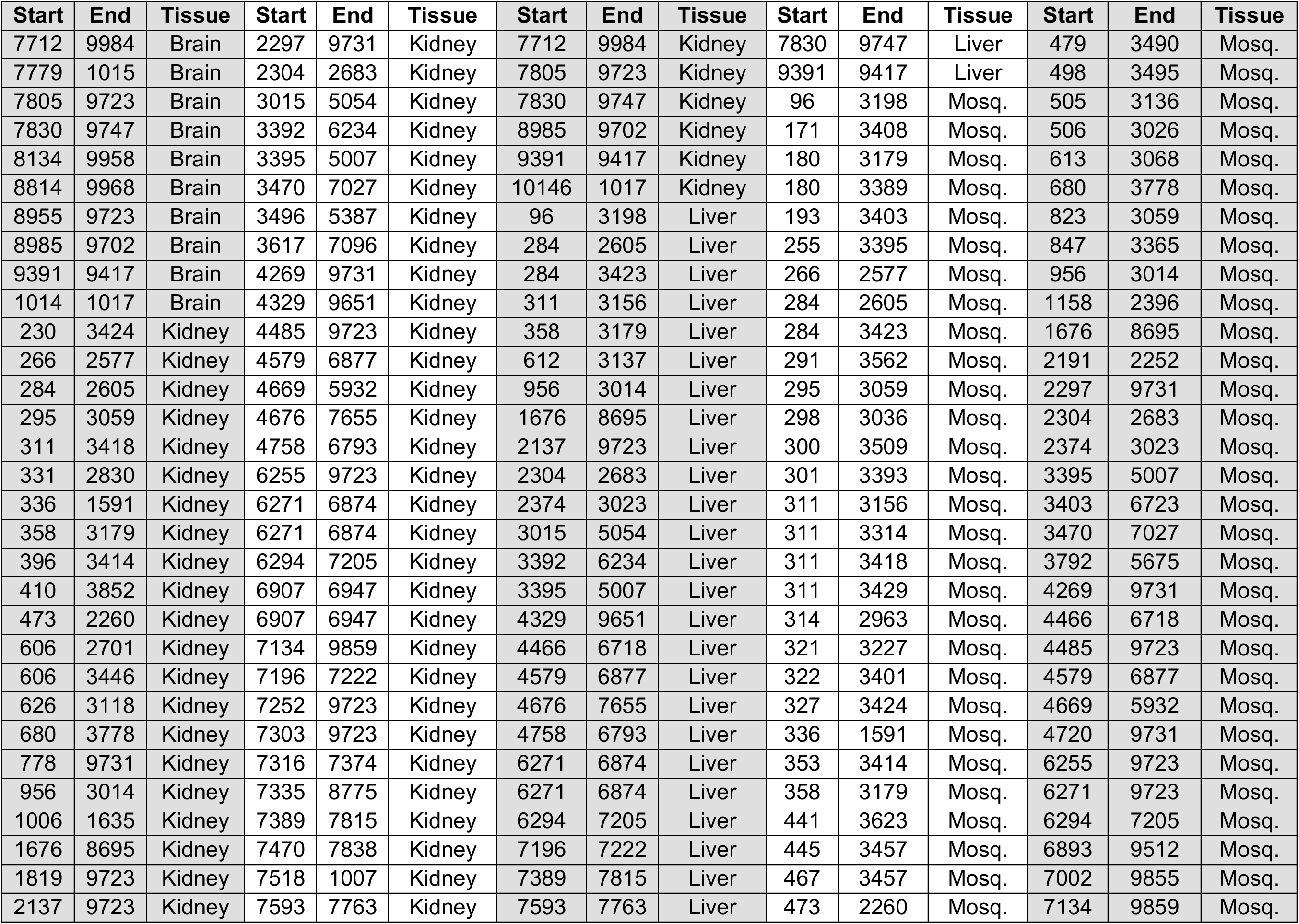
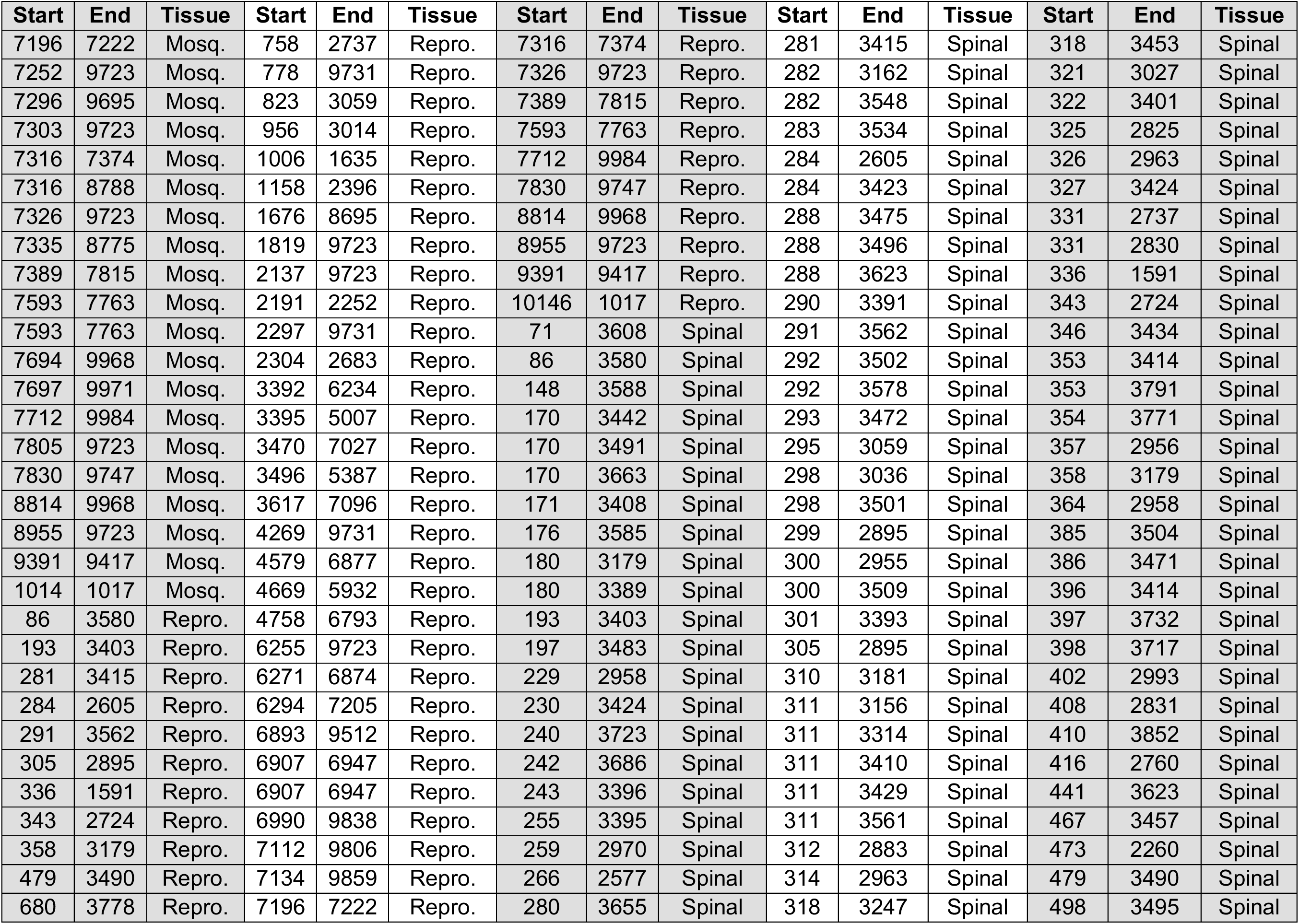
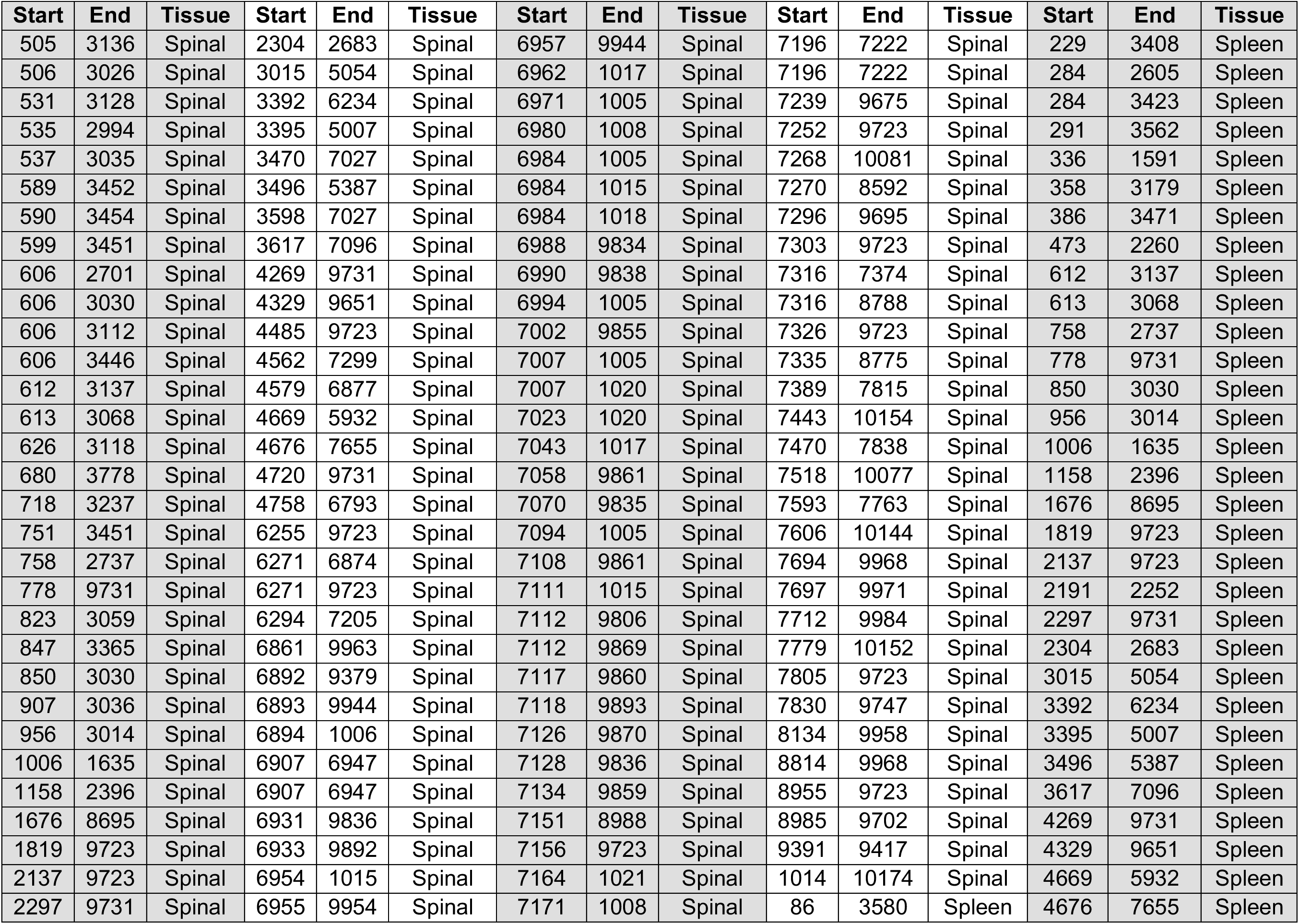
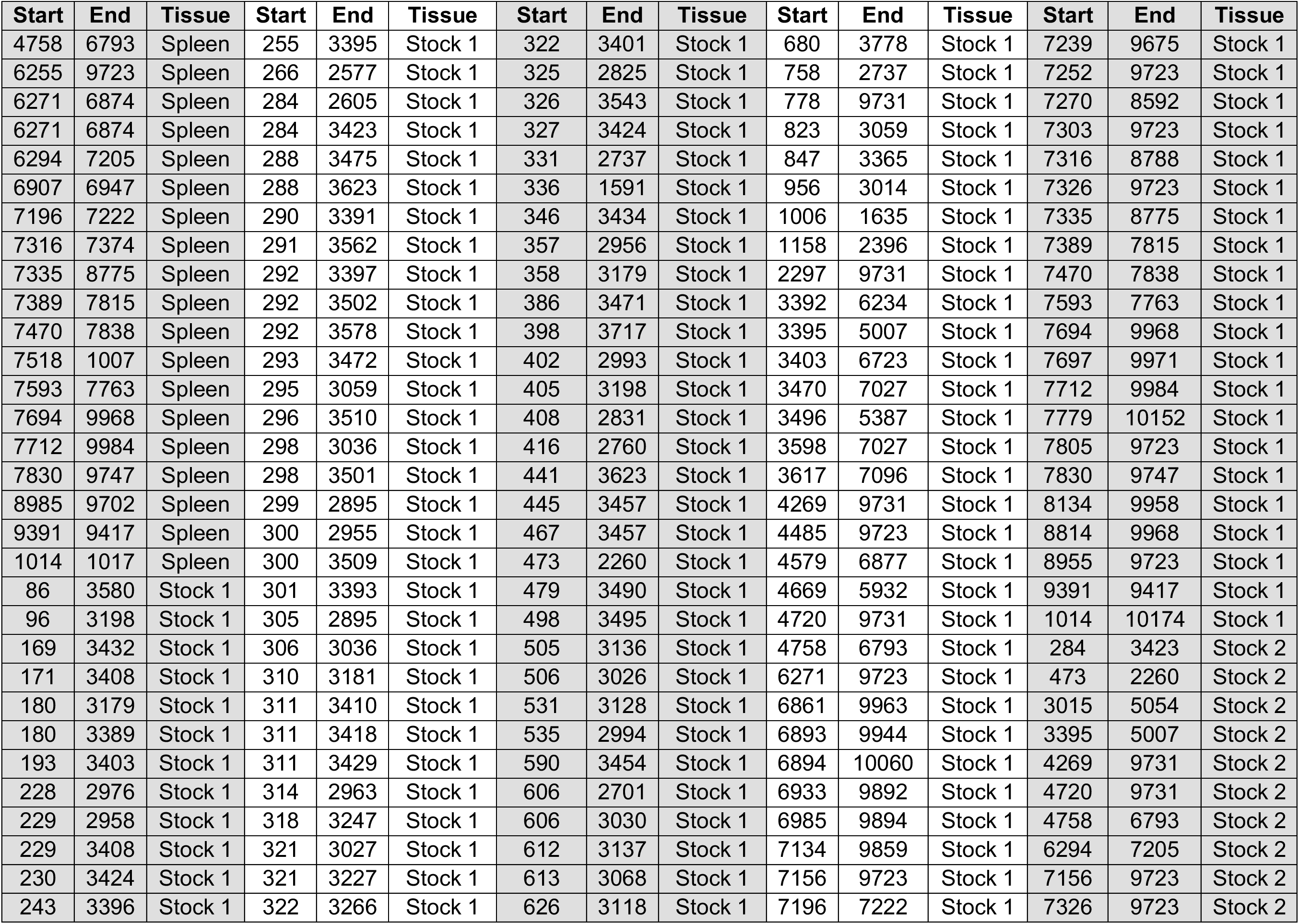
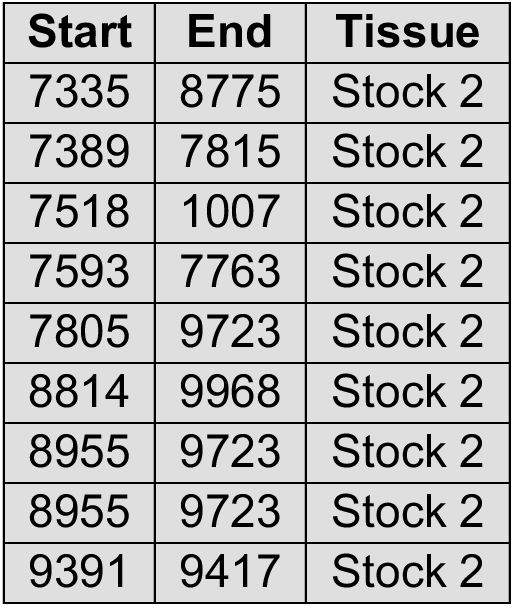
Genomic coordinates of top DVGs.

**Supplemental Table 6:**
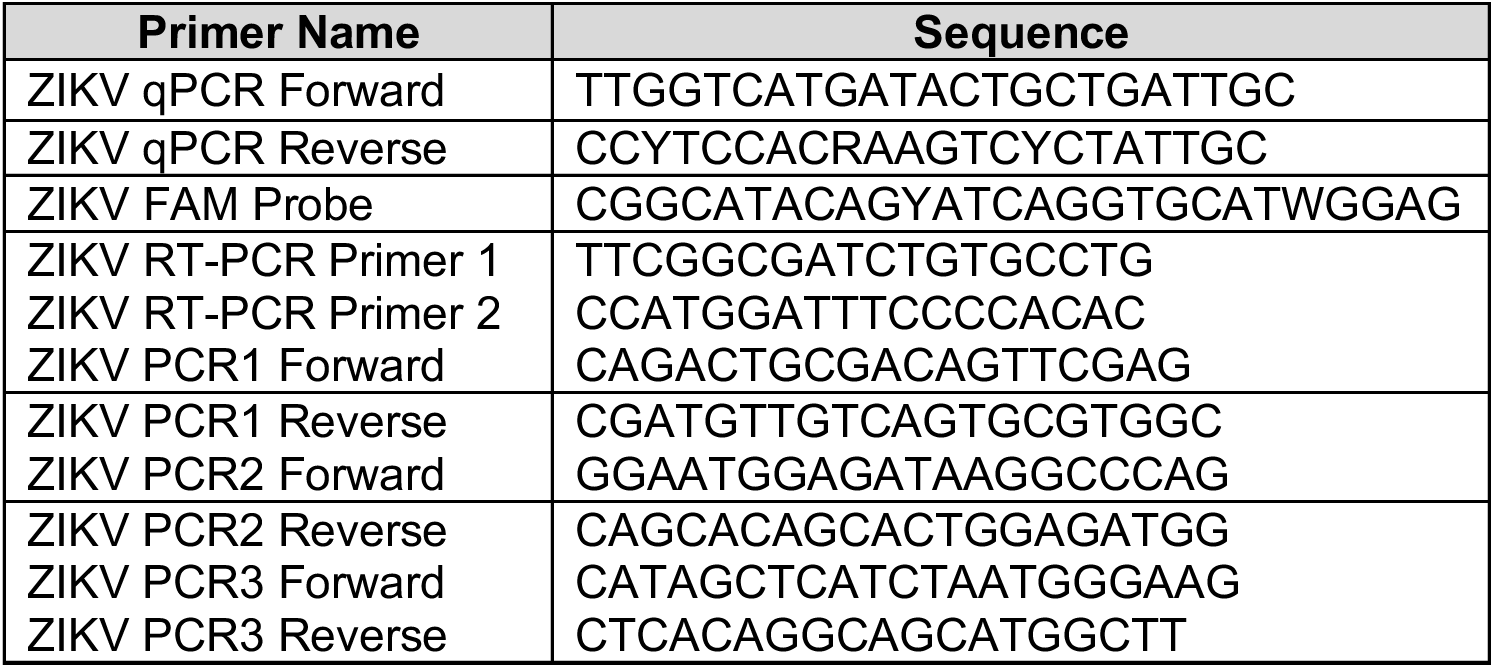
ZIKV primers used in this study

**Supplemental Figure 1:**
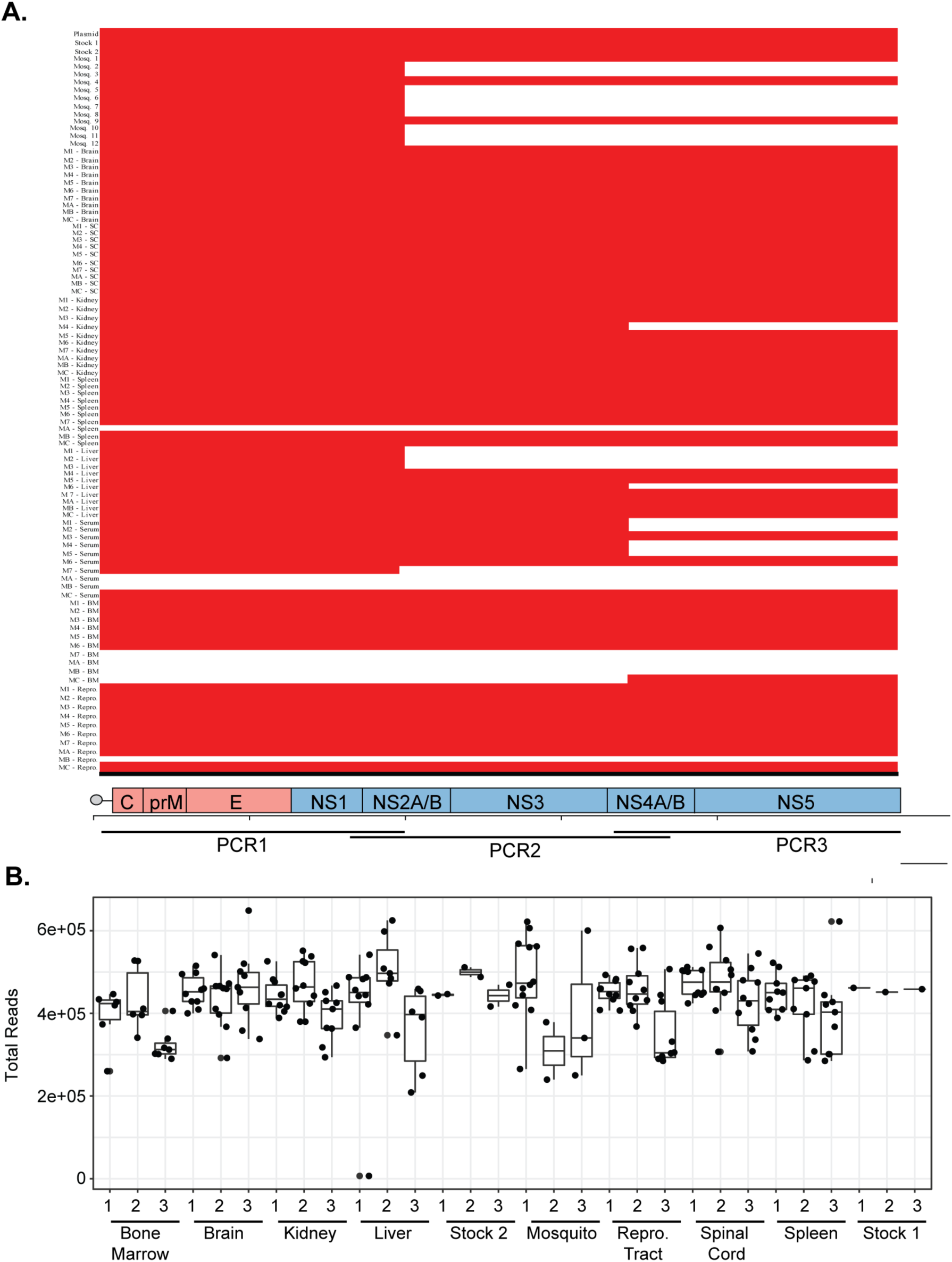
**(A).** Coding region coverage of each sample. SC = Spinal Cord. BM = Bone Marrow. **(B).** Total deep sequenclng reads of each PCR fragment per sample. Data represent the median with interquartile range.

**Supplemental Figure 2:**
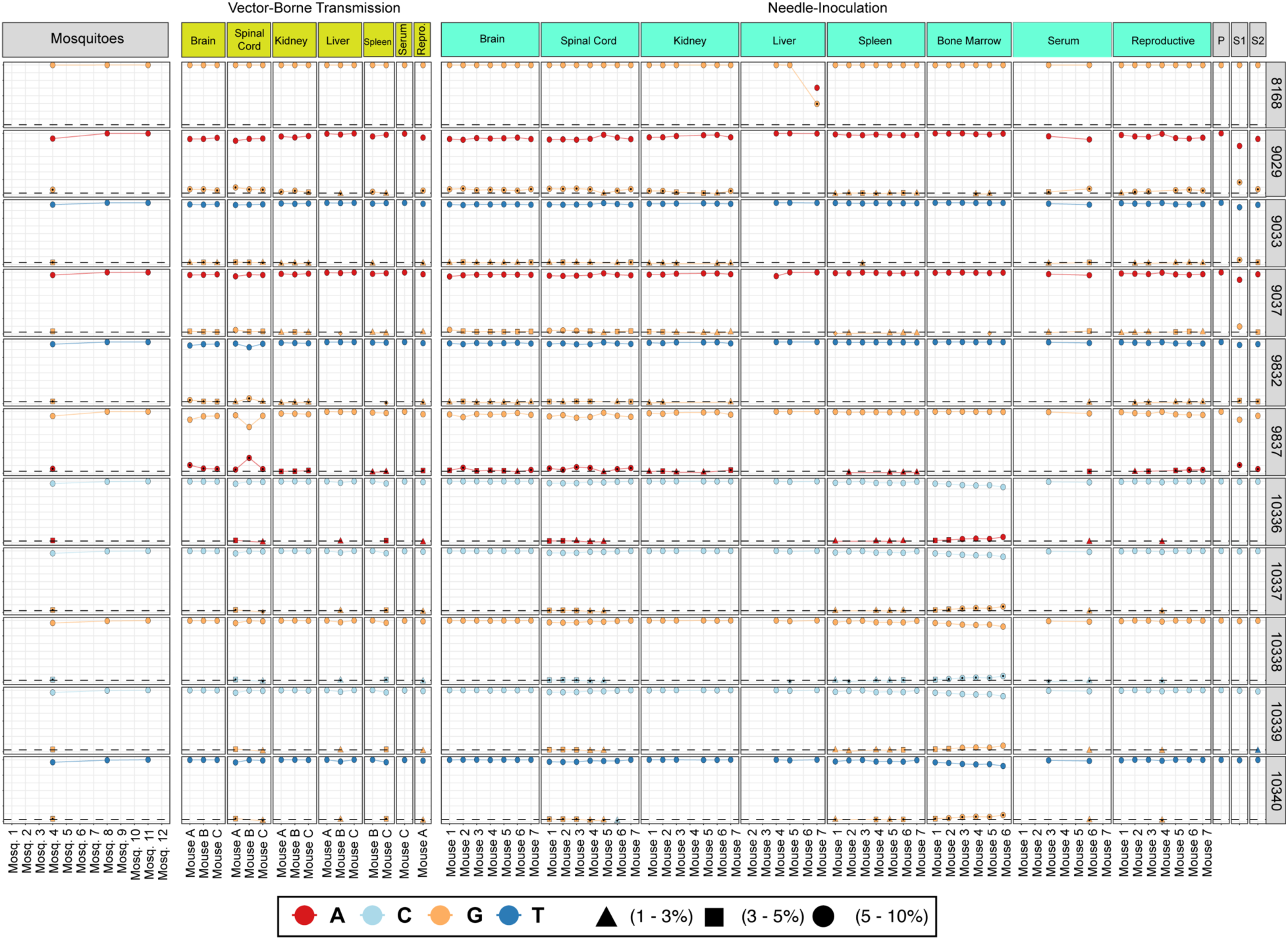
Organ-specific minority variants found in this study.

**Supplemental Figure 3:**
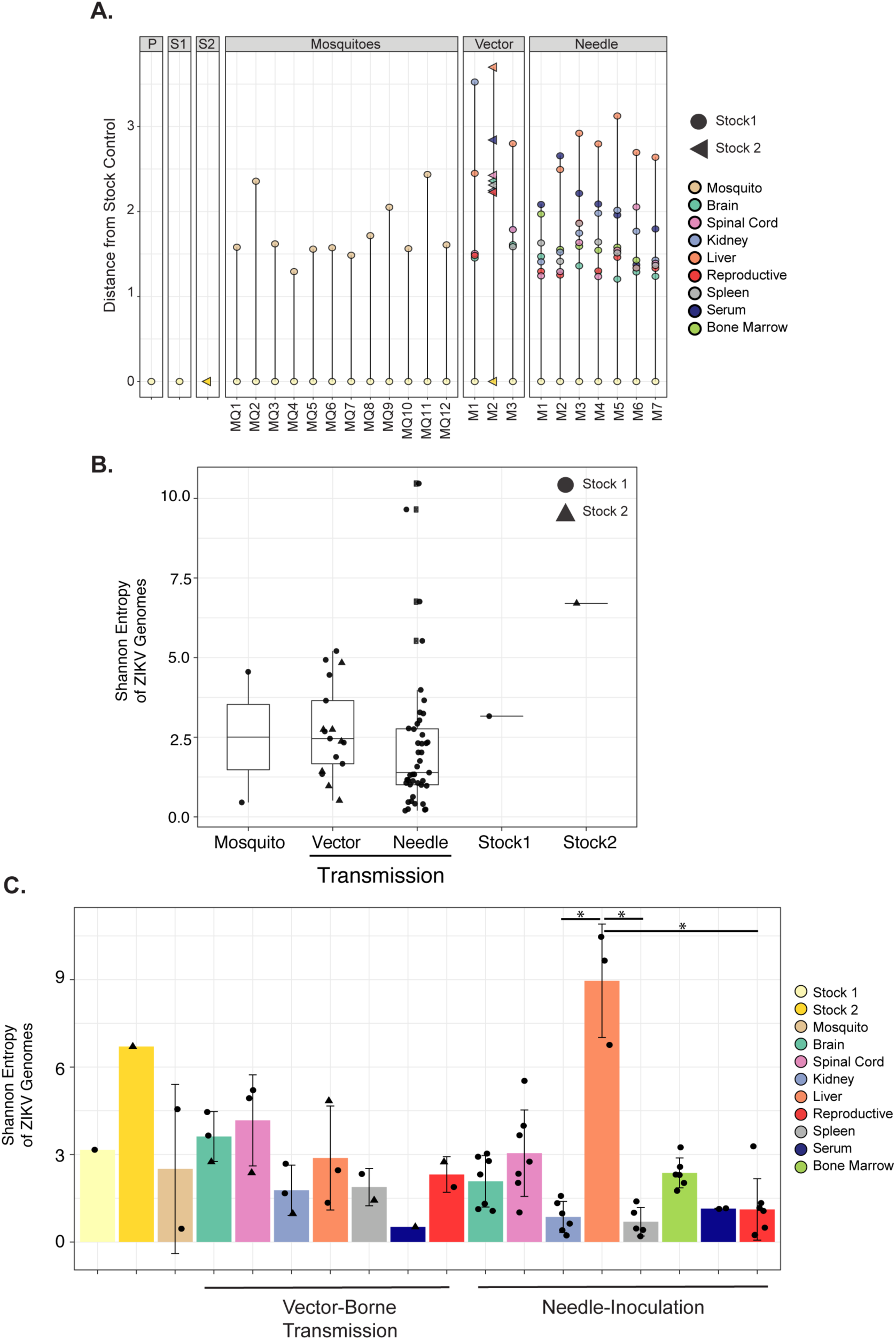
Euclidean distance per mouse and Shannon entropy over the full coding region. **(A).** Euclidean distance of each individual mosquito and mouse. **(B).** Shannon entropy by transmission route calculated over the complete coding region. **(C).** Shannon entropy of each organ per transmission route calculated over the complete coding region. Data represent the average +/- the standard deviation. Kruskal-Wallis with Dunn’s post-test, considering a type-I error.* p<0.05.

**Supplemental Figure 4:**
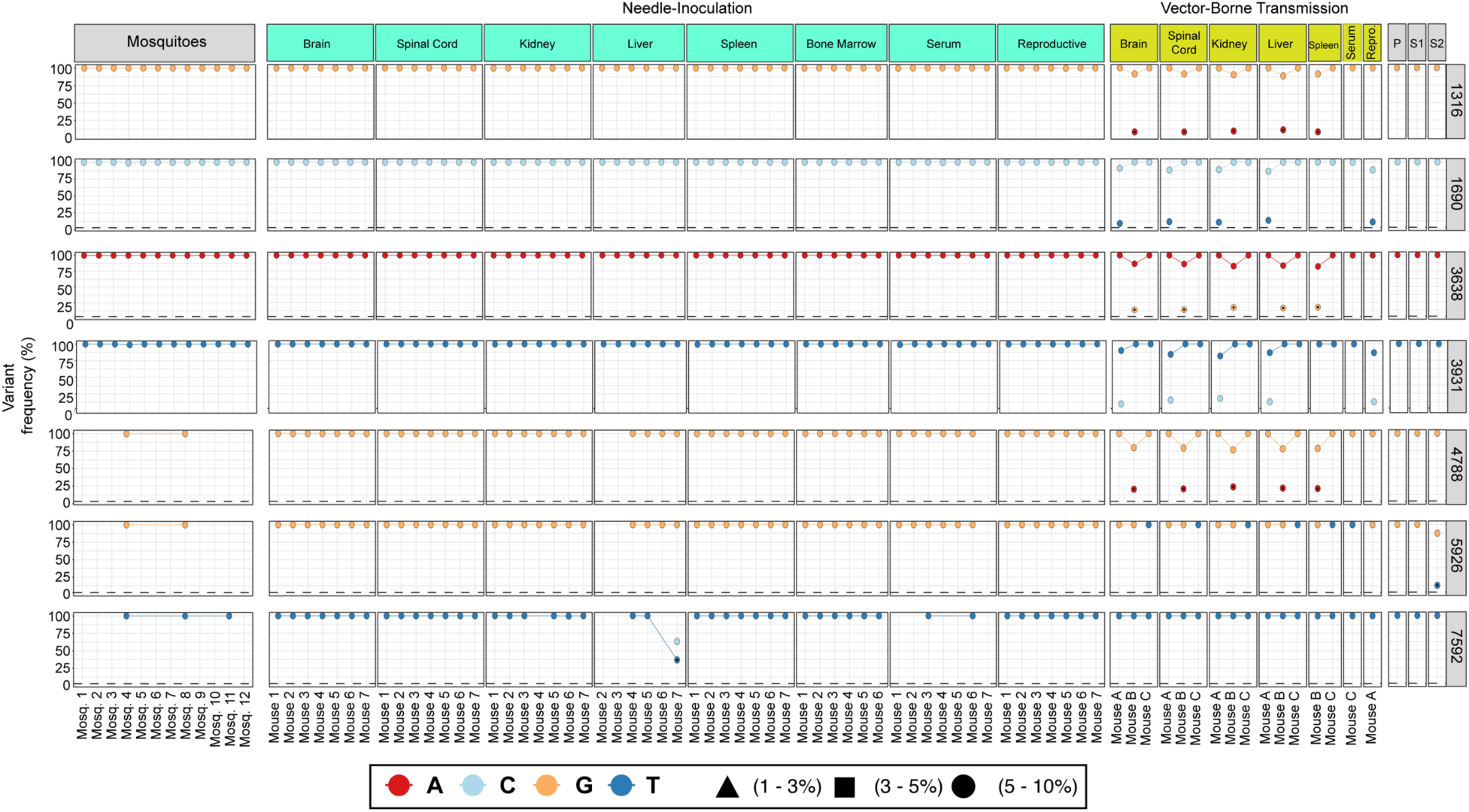
Selection of novel ZIKV during variants during vector-borne transmission.

**Supplemental Figure 5:**
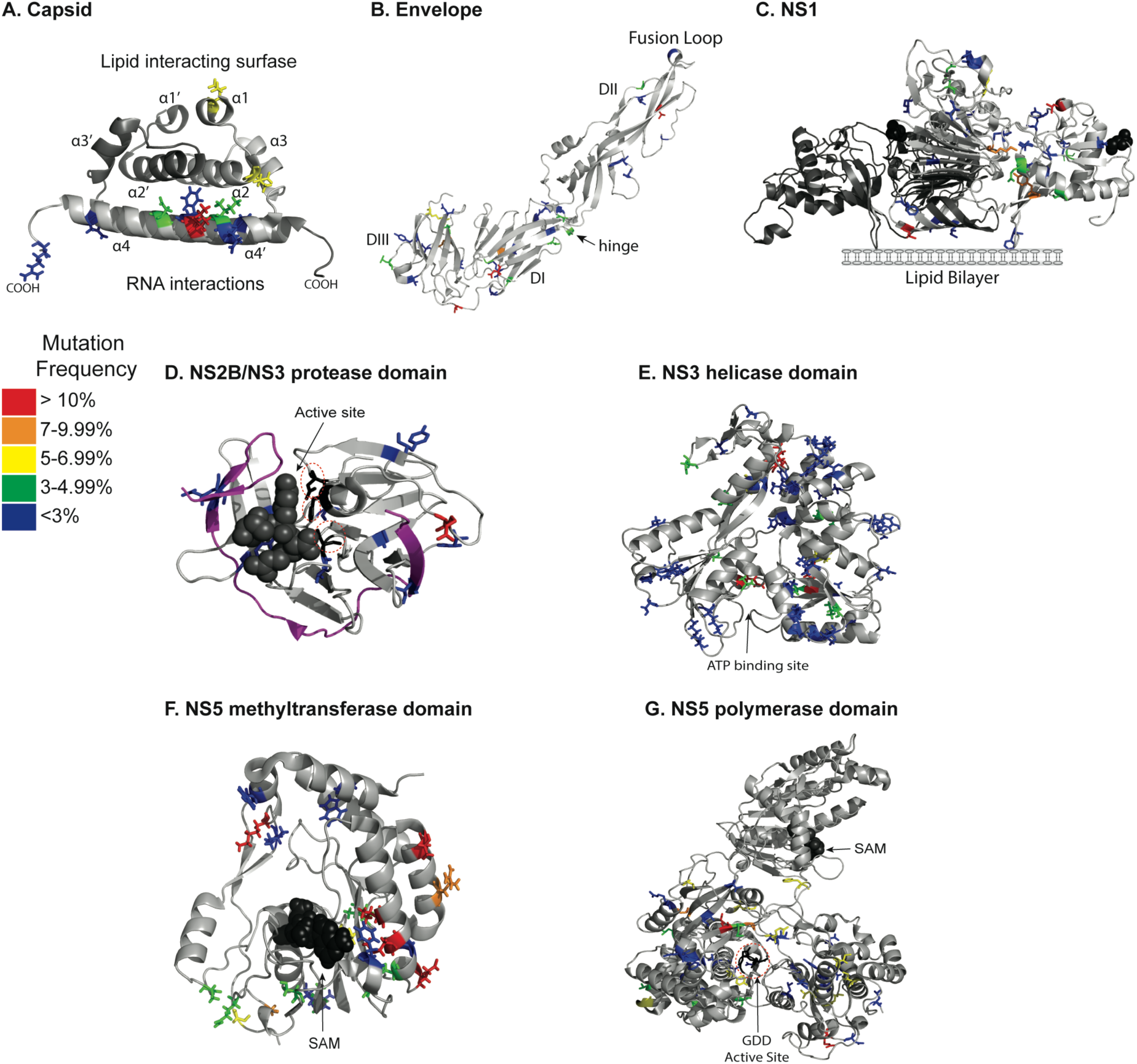
ZIKV infection highlights minority variants with potential impacts on viral biology. ZIKV minority variants were mapped onto the available protein structures of the individual ZIKV proteins (A) capsid (PDBID: 6C44), lipid- and RNA interacting surfaces are marked, (B) envelope (PDBID: 5JHM), (C) NS1 (PDBID: 5K6K), lipid bilayer indicated. (D) NS2B-NS3 protease. (PDBID: 5GJ4). NS2B is shown in magenta. Dashed circles indicate residues H51, D75, and S135 from the catalytic triad is marked in black. (E). NS3 helicase. (PDBID: 5Vl7). ATP-binding site is indicated. (F) NS5 methyltransferase domain. (PDBID: 5VIM). S-adenosyl-methionine (SAM) molecule binding site shown. (G) NS5 methyltransferase and polymerase. (PDBID: 5U0B). SAM molecule and GOD active site indicated.

**Supplemental Figure 6:**
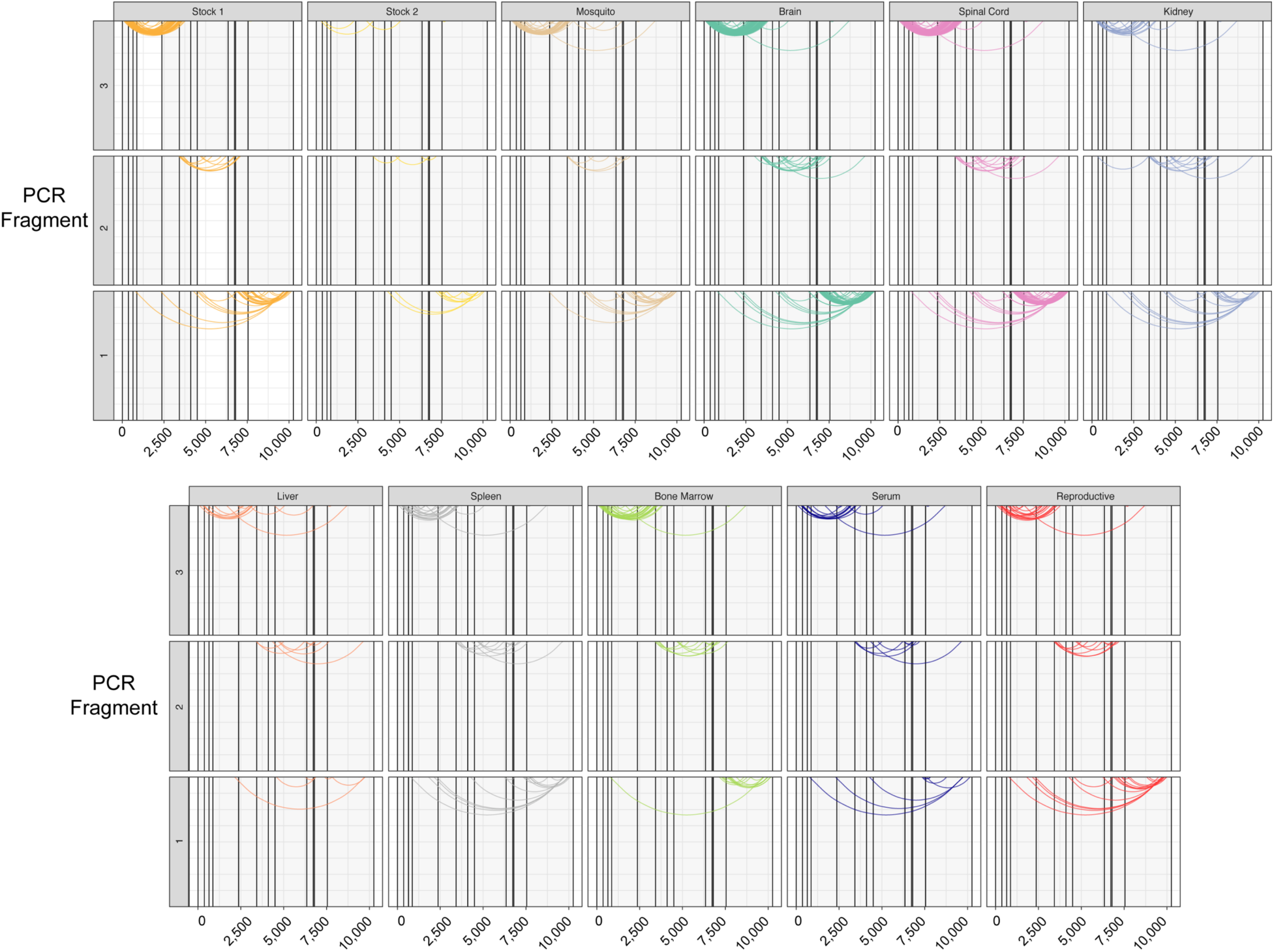
Top DVGs present in each organ. Top DVGs are defined as those found in at least 3 samples to have at least 3 reads that cover the DVG. X-axis represents nucleotide position in the ZIKV genome.

**Supplemental Figure 7:**
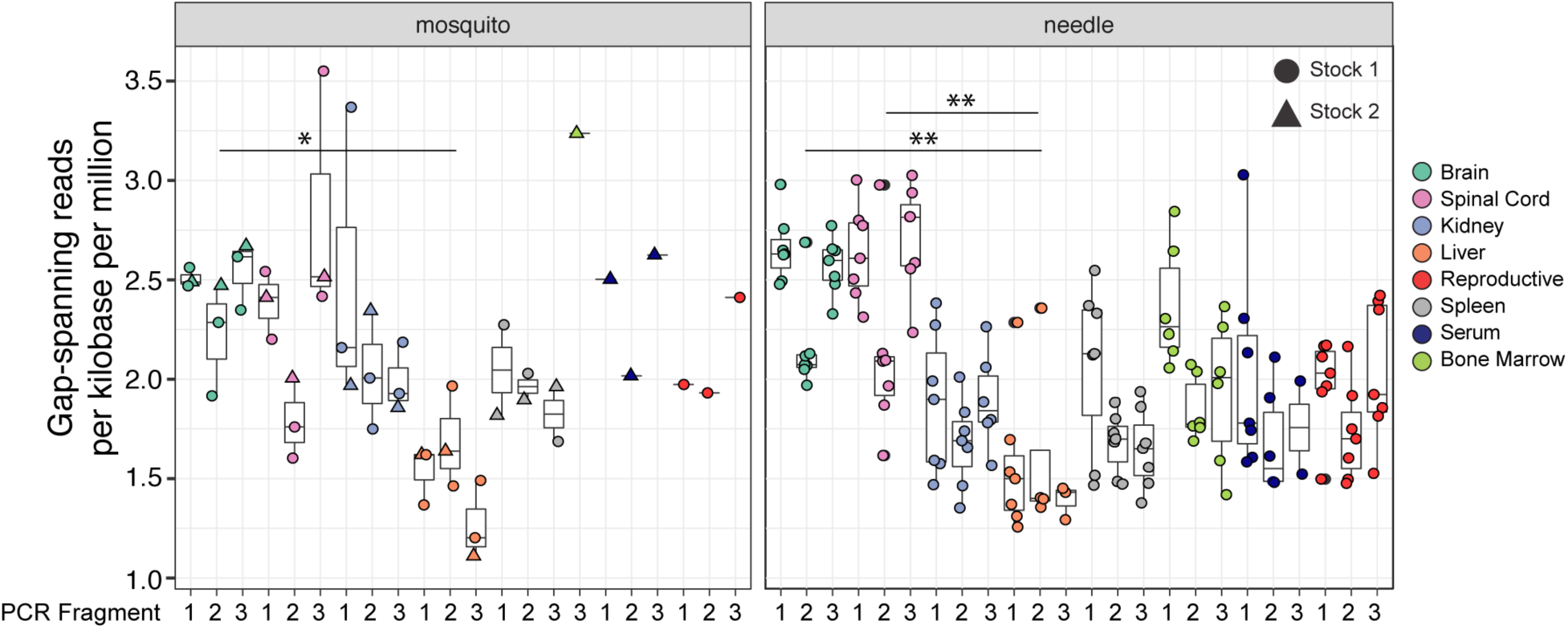
Organ-specific DVGs generated by each transmission route. Data represent the median with intraquartile range. Black dots represent outliers. Kruskal-Wallis with Dunn’s post-test, considering a type-I error. * p<0.05, ** p<0.01.

